# TMK interacting network of receptor like kinases for auxin canalization and beyond

**DOI:** 10.1101/2025.02.28.640727

**Authors:** Aline Monzer, Ewa Mazur, Lesia Rodriguez, Michelle Gallei, Minxia Zou, Michael Smejkal, Ema Červeňová, Jiří Friml

## Abstract

Receptor-like kinases (RLKs), particularly the Transmembrane Kinase (TMK) family, play essential roles in signaling and development, with TMKs being key components of auxin perception and downstream phosphorylation events. While TMKs’ involvement in auxin canalization, a process essential for vasculature formation and regeneration, has been established, nonetheless, the additional signaling and regulatory partners remain poorly understood. In this study, we identify and characterize seven leucine-rich repeat RLKs (TINT1–TINT7) as novel interactors of TMK1, revealing their diverse evolutionary, structural, and functional characteristics. Our results show that TINTs interact with TMK1 and highlight their roles in regulating various developmental processes. Majority of TINTs contributes, together with TMK1, to auxin canalization, with TINT5 linking TMK1 to other canalization component CAMEL. Beyond canalization, we also establish the role of TINT-TMK1 interactions in processes such as stomatal movement and the hypocotyl’s gravitropic response. These findings suggest that TINTs, through their interaction with TMK1, are integral components of various signaling networks, contributing to both auxin canalization and broader plant development.

## INTRODUCTION

The transition of plants to life on land brought numerous challenges, requiring adaptation to constantly changing and often harsh terrestrial conditions. As sessile organisms unable to move to evade unfavorable conditions, plants evolved sophisticated mechanisms to perceive and respond to diverse environmental signals. This adaptive pressure drove the significant expansion of the plasma membrane (PM) receptor-like kinase (RLK) family in land plants, with the model *Arabidopsis thaliana* harboring over 600 RLKs (Liu et al. 2024). These receptors enable plants to detect diverse extracellular and environmental signals and activate intracellular signaling pathways crucial for adaptive growth and development (Zhu et al. 2023).

The largest subclass of RLKs is the leucine-rich repeat (LRR) RLKs (Shiu and Bleecker 2001), distinguished by a conserved cytosolic kinase domain (KD) and a variable extracellular domain (ECD) featuring different numbers of LRRs (Liu et al. 2017). This structural variability enables them to recognize a wide range of hormonal, peptide-based or other extracellular ligands and regulate diverse biological functions (Hohmann et al. 2017). For instance, BRASSINOSTEROID INSENSITIVE 1 (BRI1) perceives brassinosteroids to regulate plant growth and development (Jiang et al. 2013), while FLAGELLIN SENSING 2 (FLS2) recognizes bacterial flagellin and triggers plant immune responses (Chinchilla et al. 2007).

Auxin is an essential plant hormone that plays a critical role in growth and development (Friml 2022). It regulates a broad spectrum of biological functions through two currently known signaling pathways (Gallei et al. 2020). The first is the canonical Transport Inhibitor Response 1/Auxin Signaling F-box (TIR1/AFB)-based pathway (Salehin et al. 2015), and the second involves the cell surface complex of the Auxin-Binding Protein 1 (ABP1)/ABP1-likes (ABLs) and Transmembrane Kinase 1 (TMK1) (Friml et al. 2022; Yu et al. 2023a). Following auxin perception, the ABP1-TMK1 complex triggers an ultrafast phosphorylation response of thousands of proteins (Friml et al. 2022; Kuhn et al. 2024), potentially regulating diverse cellular processes.

The TMK family in *Arabidopsis thaliana* includes four LRR-RLKs (Dai et al. 2013). TMK1 was first identified in 1992 (Chang et al. 1992) and later linked to key processes in auxin-regulated plant development. TMK1 regulates processes, such as root growth (Li et al. 2021), lateral root development (Huang et al. 2019), and the interdigitation of epidermal pavement cells (PCs) (Xu et al. 2014; Yu et al. 2023a). Additionally, TMK1 is implicated in apical hook maintenance (Cao et al. 2019; Gu et al. 2022; Wang et al. 2024) through a unique mechanism where its kinase domain is cleaved and translocated to the nucleus, where it phosphorylates transcriptional regulators. In other processes, such as root bending (Marquès-Bueno et al. 2021; Rodriguez et al. 2022) or organogenesis (Wang et al. 2022), TMK1 interacts with and phosphorylates PIN auxin transporters (Luschnig and Friml 2024), thereby regulating intercellular auxin fluxes. TMKs, particularly TMK4 (Wang et al. 2020; Li et al. 2024), are also involved in the regulation of auxin biosynthesis and mediate crosstalk between auxin and other hormones, such as brassinosteroids (Yu et al. 2023b) and abscisic acid (Yang et al. 2021).

A particularly fascinating and important auxin-dependent process is auxin canalization (Sachs 1981; Hajný et al. 2022). Here, the gradual formation of narrow PIN-expressing, auxin transporting channels, involving coordinated tissue polarization and specification, provides positional information for vasculature formation. Auxin canalization relies on a feedback between auxin signaling and directional auxin transport, primarily mediated by PINs (Friml 2022). This process ensures that newly formed organs integrate seamlessly with the preexisting vasculature (Benková et al. 2003; Balla et al. 2011), mediates formation (Scarpella et al. 2006) and regeneration (Mazur et al. 2016) of vasculature, thus maintaining connectivity and functional coherence within the plant. Both TMK1 and ABP1 play critical roles in auxin canalization, specifically in vasculature formation and regeneration (Friml et al. 2022). Another complex at the cell surface, involved in auxin canalization by phosphorylating PIN1, consists of two other LRR-RLKs, the Canalization-related Auxin-regulated Malectin-type RLK (CAMEL) and the Canalization-related Receptor-like Kinase (CANAR) (Hajný et al. 2020). However, auxin canalization is a complex mechanism (Hajný et al. 2022), and many additional players and regulatory mechanisms are yet to be discovered.

To gain novel insights into mechanism of canalization and TMK regulation, we identified, among the PM network of LRR-RLKs, TMK interacting partners (TINTs), which act as potential co-receptors or regulators. Here, we characterize seven of these interactors and explore their roles in auxin signaling and plant development, with a particular focus on auxin canalization.

## RESULTS

### Bioinformatical identification and characterization of TMK interactors

To identify additional players involved in TMK functions, we explored the cell surface interaction network of *Arabidopsis* (Smakowska-Luzan et al. 2018), which encompasses thousands of binary interactions between the extracellular domains of different LRR-RLKs.

Mining this dataset for TMK1-4 revealed an extensive network of TMK interactors (Fig. 1a). From this network, we selected seven most promising potential TMK1 interactors, whose interaction with TMK1 we confirmed *in planta* (Fig. S2). We designated these <u>T</u>MK <u>INT</u>eractors as TINT1 through TINT7 (Fig. S1b). Some of the TINTs have been previously mentioned in different contexts. TINT1 (also known as LRR1) is involved in plant responses to drought (Chen et al. 2021). TINT3 (Pollen-specific Receptor-like Kinase 7, PRK7) belongs to the family of pollen-specific receptor-like kinases (Takeuchi and Higashiyama 2016). TINT6 (Clavata3 Insensitive Receptor Kinase 4, CIK4) functions as a co-receptor for various receptors, prominent among them, the CLAVATA family (Hu et al. 2018; Escocard De Azevedo Manhães et al. 2021).

**Figure 1.**
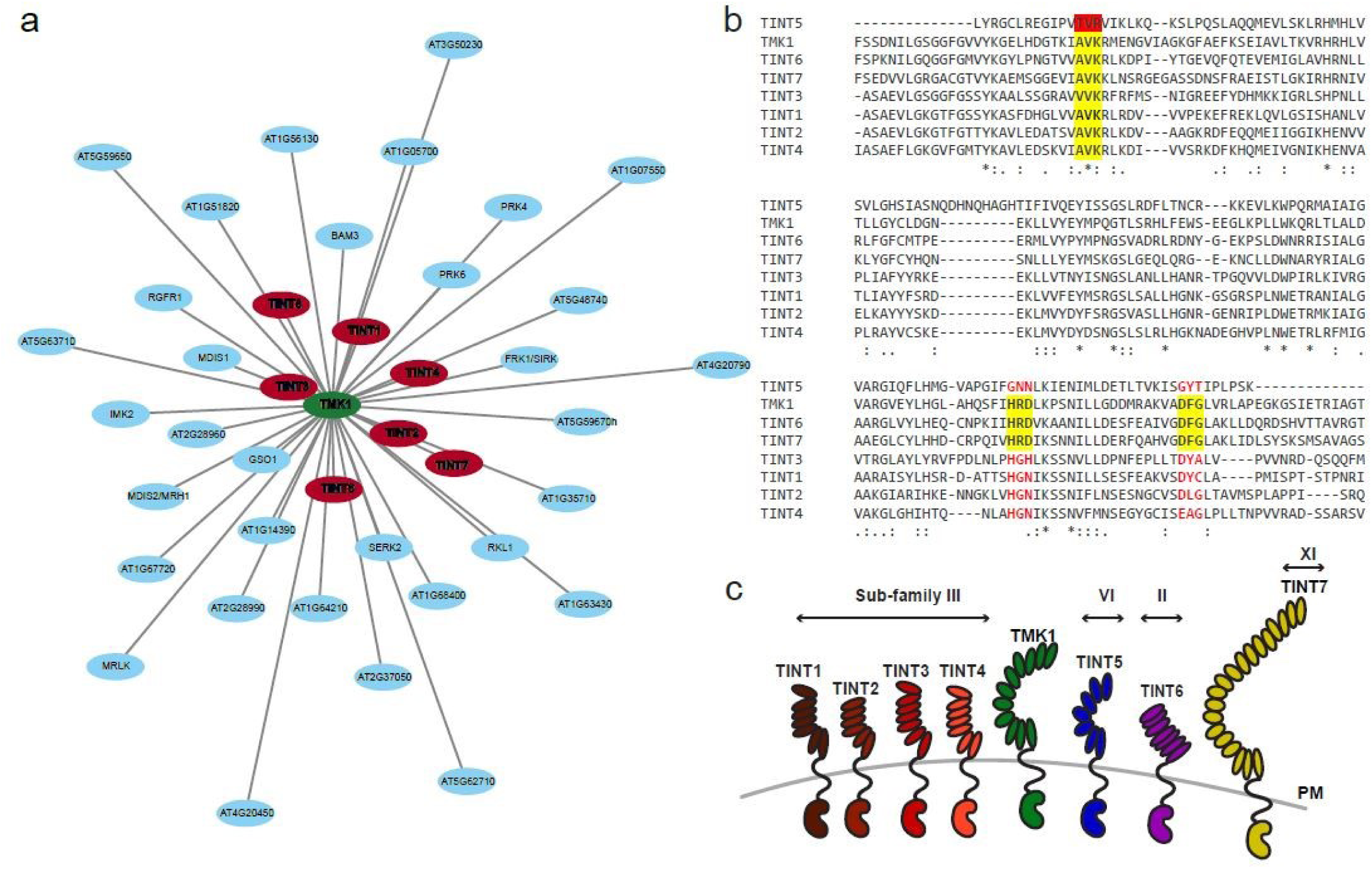
Overview of TINTs. (a) Interaction network of TMK1 (in green). Nodes represent individual proteins, and edges indicate predicted or known interactions. Key candidates, TINT1 through TINT7, are displayed in red. (b) Sequence alignment of the kinase domains (KD) of TINT1 through TINT7 and TMK1. Conserved residues are marked with asterisks (*), colons (:), and periods (.) representing different degrees of conservation across sequences. Key functional motifs, including ATP-binding lysine (K), catalytic loop (HRD motif), and activation loop (DFG motif) are highlighted. TMK1 and TINT sequences illustrate variability in conserved motifs important for kinase activity. (c) Schematic representation of TINTs with TMK1 at the plasma membrane, illustrating their classification into sub-families and relative extracellular domain (ECD) sizes.

Phylogenetic analysis revealed that TINTs are not evolutionary close to each other or to TMKs (Fig. S1a). They belong to different subfamilies of the LRR-RLK family based on the number of LRRs in their extracellular domains. TMKs are members of subfamily IX, whereas TINT1, TINT2, TINT3, and TINT4 belong to the subfamily III, TINT5 to subfamily VI, TINT6 to subfamily II, and TINT7 - the largest of the identified interactors - to subfamily XI (Fig. 1c).

Sequence alignment of the kinase domains showed that the ATP-binding lysine (K) is conserved in all TINTs except TINT5 (Fig. 1d). However, subdomain VIb (HRD motif), which forms the catalytic loop, and subdomain VII (DFG motif), which constitutes the activation loop, are conserved only in TINT6 and TINT7 and show alterations in other TINTs (Fig. 1b). This suggests that TINT1 – TINT5 may function as kinase-dead or pseudokinases.

Overall, these analyses identified seven TMK interactors from the LRR-RLKs family, constituting an evolutionary and structurally diverse group; likely including both active and non-active kinases. This diversity is consistent with the role of TMKs as central regulators of many different developmental processes.

### Confirmation of interaction between TMK1 and TINTs

We employed multiple approaches to confirm *in vivo* the interaction between TMK1 and the selected interactors.

Using a bimolecular fluorescence complementation (BiFC) assay in *Nicotiana benthamiana* leaves, we verified the interaction between the full-length TMK1 with TINT2, TINT3, and TINT4 (Fig. S2a). TMK1 dimerization served as a positive control, whereas TMK2 was used as a negative control for the interaction with TMK1.

We also performed co-immunoprecipitation (co-IP) experiments using different plant tissues. First, we conducted co-IP from *Arabidopsis thaliana* seedlings expressing HA-tagged TINT1, TINT2, TINT3, TINT4, or TINT5 under the ubiquitin (UBQ) promoter. We used an anti-HA antibody for immunoprecipitation, followed by immunoblotting with a specific anti-TMK1 antibody. The results showed that TMK1 can be co-immunoprecipitated with these interactors (Fig. S2b).

To test the interaction of TMK1 with TINT6 and TINT7, we co-expressed FLAG-tagged TMK1 with HA-tagged TINT6 or TINT7 in *Nicotiana benthamiana* leaves. Immunoprecipitation of TINT6 or TINT7 with an anti-HA antibody followed by immunoblotting with an anti-TMK1 antibody demonstrated a co-immunoprecipitation between TMK1 and these two TINTs (Fig. S2c).

Altogether, these results provided *in vivo* confirmation of interactions between TMK1 and TINT1 - TINT7.

### TINT5 localizes to PM, where it interacts with CAMEL LRR-RLK

During further exploration of the LRR-RLK interaction network (Smakowska-Luzan et al. 2018), we noticed that TINT5 is a potential interactor of CANAR, which forms complex with CAMEL and, similarly to TMKs, is involved in canalization (Hajný et al. 2020). To investigate this interaction, we used Förster resonance energy transfer combined with fluorescence-lifetime imaging microscopy (FRET-FLIM) in *Arabidopsis* protoplasts. We co-expressed GFP-tagged TINT5 with mCherry-tagged CAMEL or CANAR, and quantified the GFP lifetime. The GFP lifetime at the PM showed a significant decrease when TINT5 was co expressed with CAMEL, whereas no significant change was observed with CANAR (Fig. 2a. b). This suggests an interaction between TINT5 and CAMEL at the PM.

**Figure 2.**
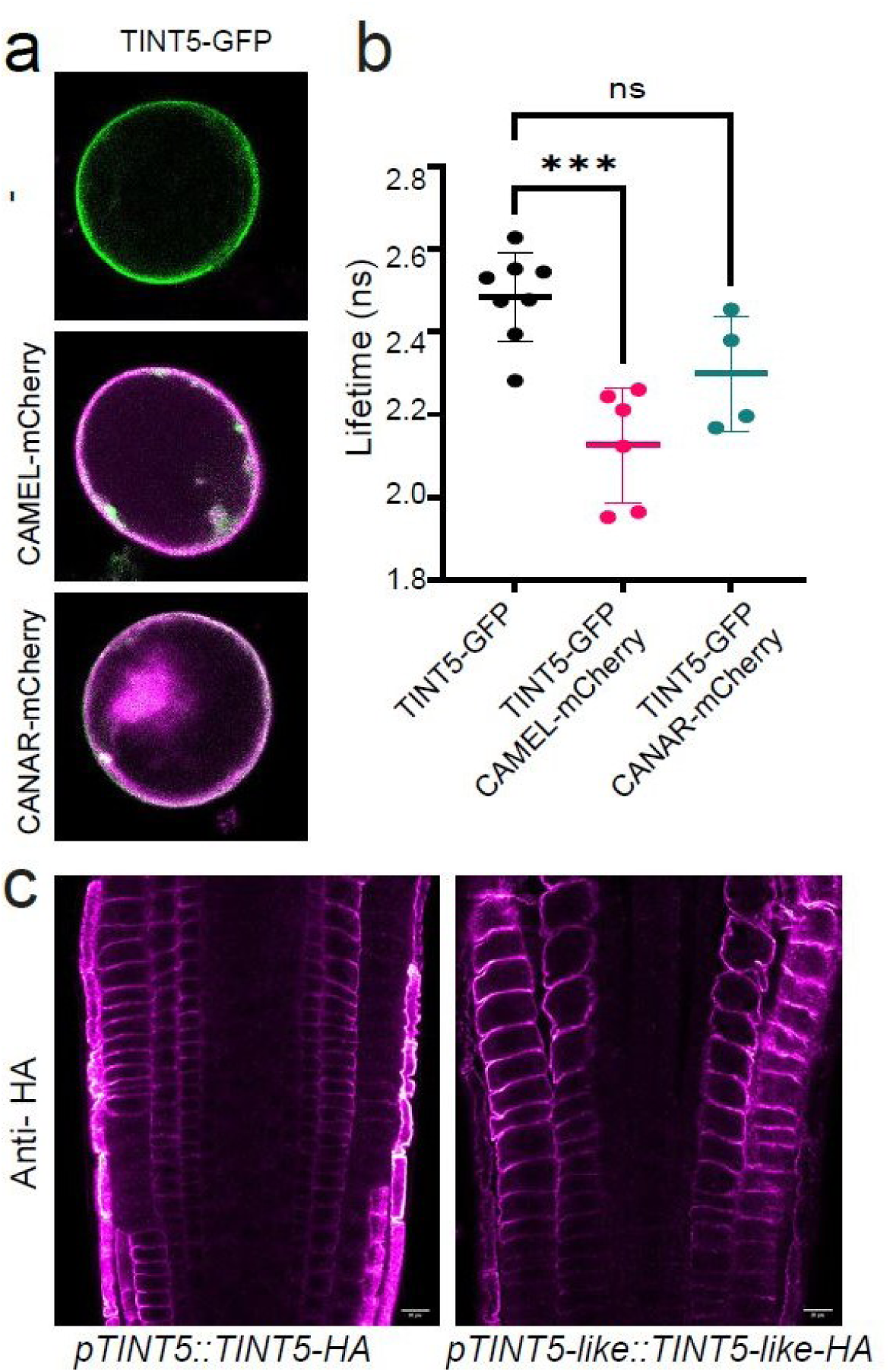
Localization of TINT5 and its interaction with CAMEL. (a) Representative images of *Arabidopsis* root protoplasts transformed with constructs encoding *p35S::TINT5-GFP* and *p35S::CAMEL/CANAR-mCherry* in FRET/FLIM assay. (b) Quantitative measurement of the GFP lifetime (FRET/FLIM) as indicated in the graph with the mean ± SD. Statistical significance is indicated by asterisks *(*** p < 0.001)* and *“ns”* for non-significant *(p > 0.05),* as determined by Dunnett’s multiple comparisons test. (c) Representative confocal images of *pTINT5::TINT5-3HA* and *pTINT5-like::TINT5-like-3HA* primary root tips after immunostaining with an HA antibody, showing the localization of TINT5 and TINT5-like. TINT5 is localized at the plasma membrane of the endodermal, epidermal and cortical cells, while TINT5-like is localized in the epidermal and cortical cells.

To confirm the PM localization, a characteristic presumably shared by all LRR-RLKs (Jose et al. 2020), we examined TINT5 as an example. We identified a homologous gene to TINT5, sharing 78% nucleotide sequence similarity, which we named TINT5-like. We generated the *pTINT5::TINT5-3HA* and *pTINT5-like::TINT5-like-3HA* transgenic lines to investigate their cellular localization by immunolocalization in *Arabidopsis* seedlings using anti-HA antibodies. This revealed that TINT5 is localized at the PM of the endodermal, epidermal and cortical cells; in the latter two cell files, it co-localizes with TINT5-like (Fig. 2c).

These observations on the example of TINT5 and its homologue show that TINTs are localized at the PM and may link TMKs to other LRR-RLK canalization components such as CAMEL/CANAR.

### *TINT*s’ expression pattern in seedlings

To gain insights into the expression pattern of *TINT* genes, we first explored publicly available transcriptome data databases (Zhang et al. 2020). The median Fragments per Kilobase of transcript per Million mapped reads (FPKM) expression values across plant tissues for *TINT1* to *TINT7* and *TMK1*, revealed distinct expression patterns among these genes (Fig. S3b). *TINT3*, *TINT4*, *TINT6*, and *TINT7* showed low to no expression in specific tissues, while *TMK1*, *TINT1*, *TINT2*, and *TINT5* exhibited moderate to high expression in several tissues.

Next, we examined their spatial expression patterns more closely using transgenic lines harboring a β-glucuronidase (*GUS*) reporter gene driven by the promoter of each *TINT* gene and compared them with *pTMK1::GUS* line in our analysis (Fig. S3a). 5-days-old seedling analysis revealed that *TINT1* is strongly expressed in the elongation zone of the root, with a weaker expression extending towards the root tip and in the vasculature of the cotyledons. *TINT2* showed more focused expression in the root tip, particularly in the root cap and meristematic zone, with some extension into the elongation zone and also in the root-hypocotyl junction. *TINT3* was primarily expressed in the root-hypocotyl junction and in lateral root primordia. *TINT4* expression was restricted to the root-hypocotyl junction. *TINT5* showed expression in both the root tip and the root-hypocotyl junction, while *TINT6* was localized to the root cap and meristematic zone in the root tip. Finally, *TINT7* was exclusively expressed in the cotyledons, showing a strong and distinct pattern along its vasculature and also in the stomata. The *pTINT1-7::GUS* expression patterns largely overlaps with that of *pTMK1::GUS*, in particular in the root tip.

As expected, the *TINT* genes show overlapping but distinct expression patterns in *Arabidopsis* seedlings. Most *TINT* genes are expressed in root regions, while *TINT7* is uniquely expressed in the cotyledon vasculature and stomata.

### *TINTs’* expression during vasculature regeneration in inflorescence stems

As the main developmental role of the ABP1-TMK auxin perception complex is in auxin canalization-mediated vasculature regeneration (Friml et al. 2022), we focused on the role of *TINTs* in this process. First, we analyzed their spatial and temporal expression patterns in inflorescence stems following at 2, 4 and 6 days after wounding (DAW), which interrupted the stem vasculature. Using the *TINT::GUS* lines, we inferred TINT promotor activity associated with vasculature regeneration (Fig. 3).

**Figure 3.**
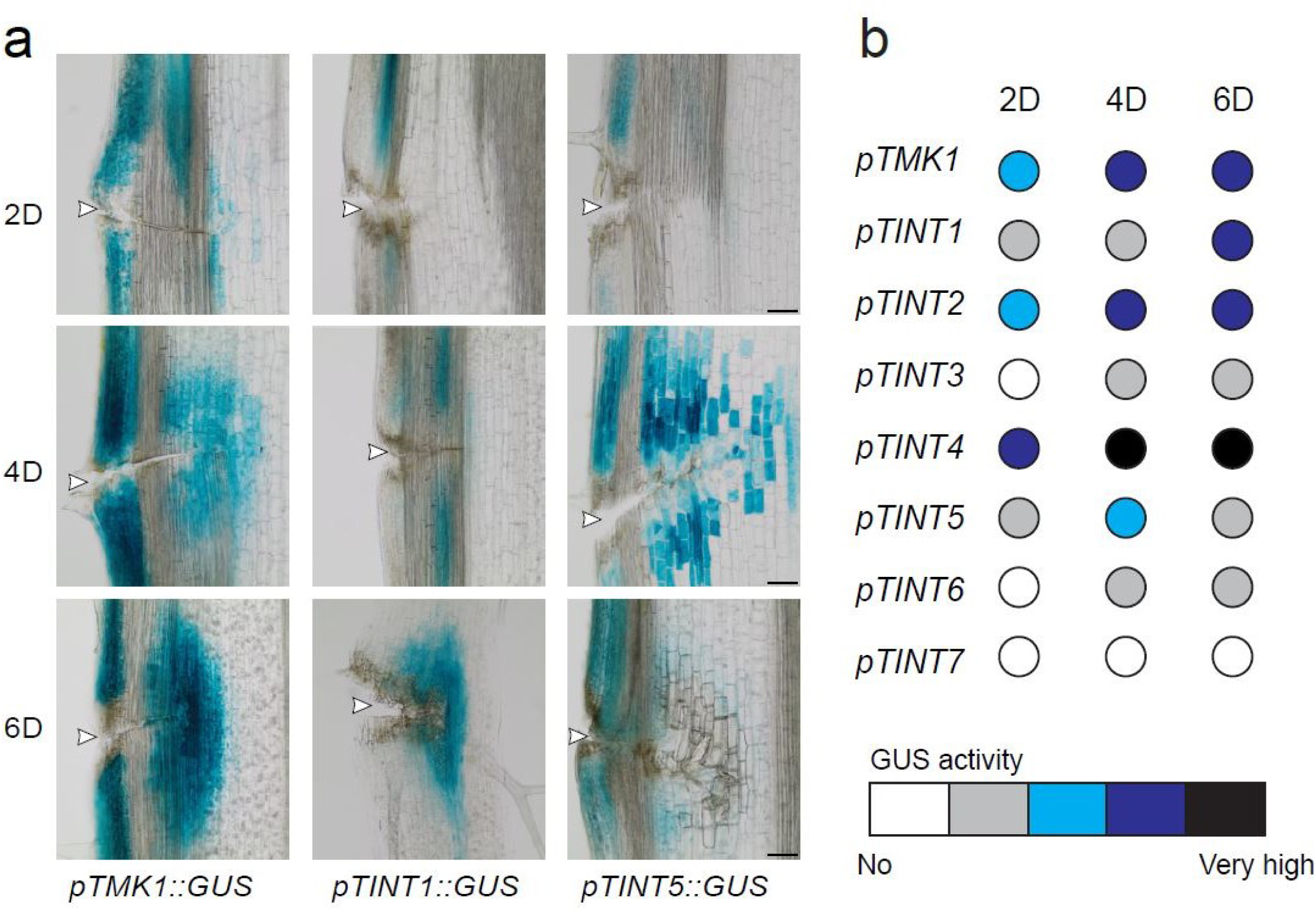
*TINT* expression during vasculature regeneration. (a) Representative images of the GUS staining of inflorescence stems during vasculature regeneration at 2, 4 and 6 days after wounding (DAW). *TMK1*, *TINT1*, and *TINT5* showed different expression patterns. White arrowheads indicate the wounding site. Scale bar, 100 µm. (b) Visual quantification of GUS staining intensity in cells surrounding the wound for each genotype at 2, 4, and 6 DAW. Color gradients represent expression levels, from white indicating the lack of expression to black indicating very high expression. The highest GUS reaction was observed in *pTINT4::GUS*. A high and extended GUS activity was also found in *pTMK1::GUS* and *pTINT2::GUS.* No *TINT7* expression was observed in the regenerated vasculature.

As shown previously (Friml et al. 2022), *TMK1::GUS* showed elevated GUS activity at 2 DAW in the outer tissues of the wounded area, with a further increase at 4 and 6 DAW, particularly around the wound (Fig. 3a). The expression was notably stronger in the differentiating cells. Similarly, *TINT1::GUS* exhibited a gradual increase in signal around the wound, with the highest levels observed at 6 DAW. *TINT2* displayed elevated expression above the wound at 2 DAW, extending around the wound at 4 and 6 DAW (Fig. S4). In contrast, *TINT3* had no expression at 2 DAW, confined to the outer tissues, with a slight extension to the area around the wound at 4 and 6 DAW. *TINT4* was expressed around and below the wound at 2 DAW, as well as in a narrow area around the wound. At 4 and 6 DAW, GUS activity was very high and extended throughout the area surrounding the wound. *TINT5* showed increased expression in the outer tissues above the wound at 2 DAW, followed by a stronger response around the wound at 4 DAW. However, by 6 DAW, *TINT5* expression decreased, remaining only in cells around the wound. *TINT6* demonstrated extended expression in cells around the wound but no activity was detected in the regenerated vasculature. *TINT7* was expressed only in the preexisting vasculature, particularly in the vascular bundles and below the wound, with no expression observed around the wound (Fig. S4).

Taken together, most of the *TINT* genes are expressed during vasculature regeneration, displaying partially overlapping temporal and spatial expression patterns with strong expression around the wound (such as *TINT1*, *TINT2* and *TINT4*), while *TINT7* remains confined to the preexisting vasculature.

### Vasculature regeneration defects in *tint* mutant stems

To test the functional requirement of TINTs in vasculature regeneration, we isolated and confirmed *tint* loss-of-function mutants. (Fig. S1c).

We assessed the ability of these mutants to regenerate vasculature in the inflorescence stems following wounding. A precise cut was made in the stems, and after 6 days, Toluidine Blue O (TBO) staining was used to visualize the differentiated vasculature (Fig.4a). In the wild type (WT) plants, fully developed vasculature successfully circumvented the wound site with visible connections between the preexisting and newly formed vasculature (Fig. 4b). Similarly, *tint1*, *tint2*, and *tint4* showed complete regeneration of vasculature around the wound (Fig. 4c, Fig. S5). In contrast, regeneration in *tint3*, *tin5*, *tint6*, and *tint7* was either partial or completely absent, with respective percentages (90%, 60%, 100% and 100%) indicating the combined frequency of partial or no regeneration observed (Fig. 4c). These vasculature regeneration defects closely resembled those observed in the *tmk* mutants (Fig. 4b). Notably, when we examined the venation patterns in the cotyledons of *tint* loss-of-function mutants, all of them showed normal development, forming the typical 4-loop vein pattern as in WT (Fig. S5c). This is consistent with largely normal venation in *tmk* mutants. Thus, *tint3*, *tint5*, *tint6* and *tint7*, similar to *tmk* mutants, show specific defects in vasculature regeneration (Fig. 4b-c).

**Figure 4.**
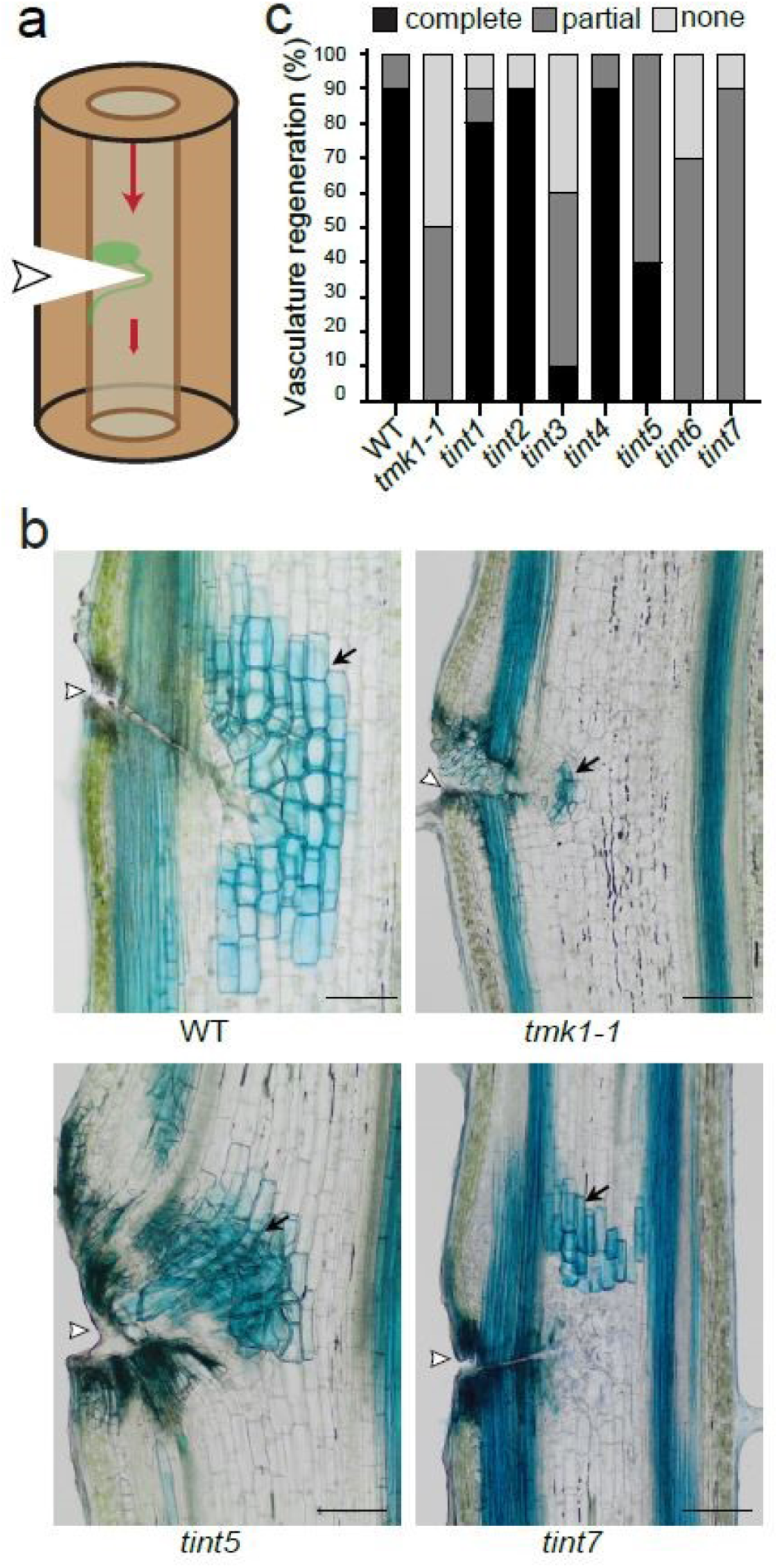
TINT and TMK1 in vasculature regeneration after wounding. (a) Schematic representation of vasculature regeneration in an inflorescence stem after wounding. White arrowhead indicates the wounding site. Red arrows indicate auxin flow. Green indicates the accumulation of auxin above the wound and the formed auxin channel guiding regenerated vasculature circumventing the wound. (b) Representative images of the TBO staining at 6 DAW showing vasculature regeneration in the inflorescence stems. WT vasculature regenerated completely around the wound, however *tmk1-1*, *tint5* and *tint7* mutants regenerated only partially. White arrowheads indicate the wounding site. Black arrows indicate the regenerated vasculature. Scale bar, 100 µm. (c) Quantification of the vasculature regeneration across genotypes, categorized as complete (fully formed vasculature), partial (limited and partially developed vasculature), or none (no vessel formed around the wound), at 6 DAW. n=10 plants per genotype.

To further test the common action of TINTs and TMKs in vasculature regeneration, we generated and tested some *tint tmk1* double mutants. The *tint5 tmk1-1* double mutant showed a more pronounced defect than the single mutants (Fig. S7 a-b). A similar trend was observed in *tint6 tmk1-1* (Fig. S5 a-b) and *tint7 tmk1-1* double mutants (Fig. S5 a-b). These observations suggest that the combined loss of TMK1 with TINT5, TINT6, or TINT7 leads to a more significant disruption in vasculature regeneration.

Altogether, these results collectively show the common function of TMK1 with TINT3, TINT5, TINT6, and TINT7 in vasculature regeneration processes, providing additional genetic support for their interaction with TMK1.

### Roles of TINT5 and TINT7 in vasculature regeneration and auxin canalization

Due to the strong defects in vascular regeneration observed in the *tint5*, *tint6* and *tint7* mutants, we selected these genes for an in-depth analysis of their role in auxin canalization.

The key feature of auxin canalization is auxin’s ability to promote and form its own transport channels, which in turn establish new vasculature paths (Hajný et al. 2022). To examine this, we applied a droplet of exogenous auxin (Indole-3-Acetic Acid, IAA) as a localized auxin source to the side of the stem below the wound, to induce *de novo* vasculature formation (Mazur et al. 2020b) (Fig. 5a). The forming vasculature was visualized using TBO staining 4 days after application (Fig. 5b). Ninety percent of WT stems formed continuous vasculature from the local auxin source; however, *tint5* and *tint6* exhibited, similarly to the *tmk1-1* mutant, reduced *de novo* vasculature formation, with this effect being more pronounced in the double mutants of *tint5 tmk1-1* and *tint6 tmk1-1* (Fig. 5c). The *tint7* and *tint7 tmk1-1* mutants showed even stronger defects; being unable to form any vasculature from the auxin source (Fig. 5c, Fig. S6a.). These results demonstrate that TINT5, TINT6 and TINT7 share a common function with TMK1 not only in vasculature regeneration but also in its *de novo* formation from the localized auxin source.

**Figure 5.**
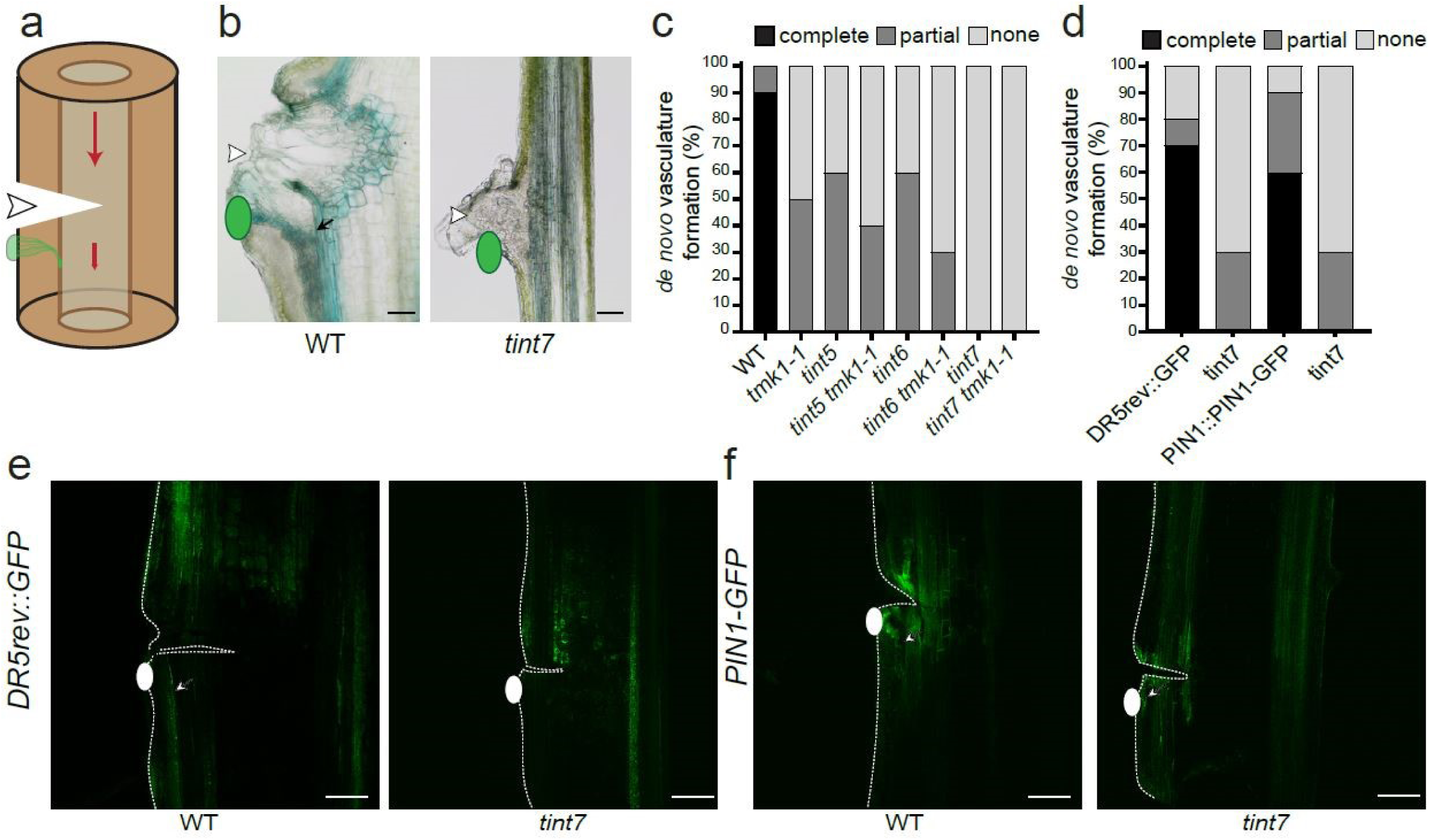
TINT and TMK1 in auxin channel formation. (a) Schematic representation of canalization and *de novo* vasculature formation from local application of auxin to wounded stems. White arrowhead indicates the wounding site. Red arrows indicate auxin flow. Green indicates the local application of auxin (IAA) and *de novo* formed auxin channel guiding formation of vasculature. (b) Representative images of the TBO staining of *de novo* vasculature formation and canalization at 6 days after IAA application (DAA) (green ovals) following wounding. WT vasculature regenerated fully from auxin application, while *tint7* showed no regeneration. White arrowheads indicate the wounding site. Black arrow indicate the regenerated vasculature from the applied source of auxin. Scale bar, 100 µm. (c) Quantification of *de novo* vasculature formation from a local source of auxin in WT, *tmk1-1*, *tint* and double mutants, categorized as complete (fully formed vasculature), partial (limited and partially developed vasculature), or none (no vessel formed) at 6 DAA. n=10 plants per genotype. (d) Quantification of *de novo* vasculature formation from a local source of auxin in *DR5rev::GFP* and *PIN1::PIN1-GFP* lines, categorized as complete (fully formed channels), partial (limited or partially developed channels), and none (no channels formed) at 4 DAA. n=10 plants per genotype. (e-f) Exogenous application of IAA (white ovals) below the wounding site to the stem of *DR5rev::GFP* (e) and *PIN1::PIN1-GFP* (f) triggered the complete development of auxin and PIN1 channels respectively (indicated by white arrows), but not in the *tint7* mutant. 4 DAA. Scale bar, 100 µm.

Auxin canalization relies on the feedback loop between auxin perception and its directional transport mediated by PIN auxin transporters, leading to a formation of PIN1-expressing auxin channels (Hajný et al. 2022). To gain insights into TINT5 and TINT7 involvement in these processes, we visualized auxin-transporting channels using the auxin response marker *DR5rev::GFP* (Friml et al. 2003) and the *pPIN1::PIN1-GFP* (Benková et al. 2003). The gradually narrowing channel-like patterns can be observed both during regeneration after wounding and during *de novo* vasculature formation from the localized auxin source (Mazur et al. 2020b; Hajný et al. 2022). Indeed, as reported in the WT, we observed the formation of DR5- and PIN1-positive channels circumventing the wound and reconnecting the preexisting vasculature (Fig. S6b). Similar DR5- and PIN1-positive channels formed from the places of local auxin application, predicting the formation of new vasculature (Fig. 5e-f). The *tint5* and *tint7* mutants exhibited defective formation of DR5- and PIN1-marked channels in both cases as compared to WT patterns (Fig. 5d, Fig. S6c)). Notably, in the mutants, the GFP-positive cells were positioned at the outer edge of the stem without forming the characteristic channel-like pattern (Fig. S6b).

Altogether, these results collectively establish the involvement of TINT5 and TINT7 in auxin channel formation in a similar way to what has been shown for TMK proteins (Friml et al. 2022).

### TINT5 and TINT5-like are redundantly involved in auxin canalization

Given that TINT5 has a close homologue, TINT5-like, we tested a potential functional redundancy between these two proteins. We crossed the *tint5* and *tint5-like* single mutants to generate *tint5 tint5-like* double mutant.

We assessed the phenotype of the *tint5 tint5-like* double mutant in vasculature regeneration after wounding and in *de novo* regeneration. Indeed, the *tint5 tint5-like* double mutant exhibited a more severe phenotype than the single mutants in both processes (Fig. S7). This supports the involvement of TINT5 and its homologue in auxin canalization.

These results confirm redundant action of TINT5 and TINT5-like in auxin canalization.

### TINT6 in auxin canalization and hypocotyl bending termination processes

Auxin-induced repolarization of PIN auxin transporters is a key component of the canalization mechanism (Hajný et al. 2022). Nonetheless, a similar phenomenon has been observed during other processes as well. For example, during shoot gravitropism, auxin-induced repolarization of PIN3 is essential for the termination of gravitropic bending (Rakusová et al. 2016). Thus, these two processes share a common feature of the auxin effect on PIN polarity.

While analyzing gravitropic response of etiolated hypocotyls in different *tint* mutants, we observed impaired responses in the *tint6* and *tint6 tmk1-1* mutants (Fig. 6a). The *tint6* hypocotyls exhibited hyperbending with an average bending angle of 56°, as compared to WT hypocotyls (angle of about 40°), while the double mutant *tint6 tmk1-1* showed reduced bending with an angle of 34° (Fig. 6b).

**Figure 6.**
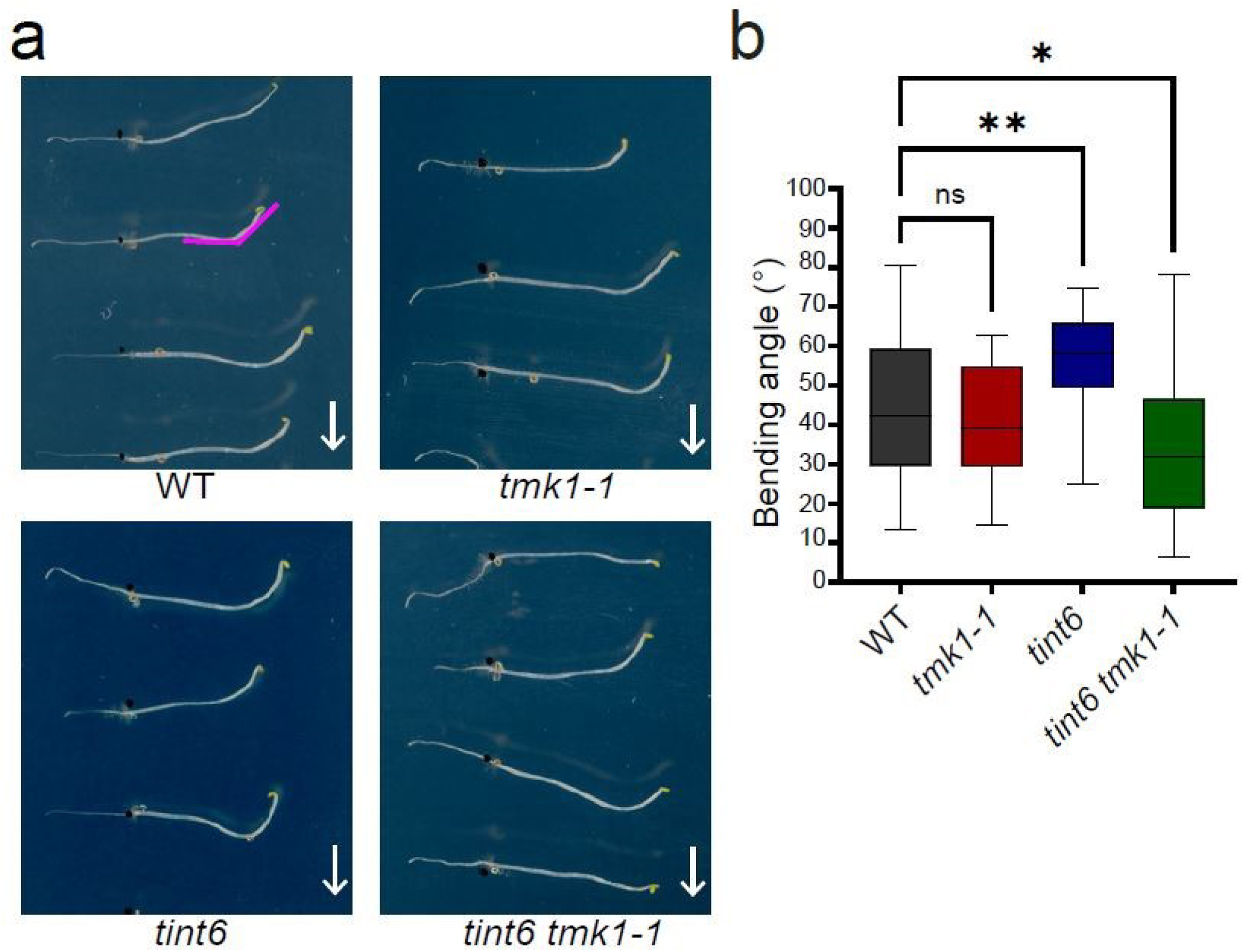
TINT6 function in regulating hypocotyl bending. (a) Representative images of the hypocotyl gravitropic response in WT, *tmk1-1*, *tint6* and the double mutant *tint6 tmk1-1* after 24 hours of gravistimulation. The white arrow indicates the direction of gravity, and the pink angle marks the curvature measured after 24 hours of gravistimulation. (b) Quantification of hypocotyl bending angles after 24 hours. Values represent the average curvature with minimum and maximum ranges. WT hypocotyls showed an average bending of 44°, *tint6* exhibited hyperbending, while *tint6 tmk1-1* showed hypobending. Statistical significance is indicated by asterisks *(* p < 0.05, ** p < 0.01) and “ns” for non-significant (p > 0.05),* as determined by Dunnett’s multiple comparisons test.

These results on the *tint6* hypocotyl bending phenotype suggest a genetic link between auxin-induced PIN repolarization during canalization and the termination of gravitropic bending.

### Roles of TINT7 and TMK1 in stomata movement

Given that *TINT7* is unique among *TINTs* for being highly expressed in stomatal cells (Fig. 7a), we investigated its potential function in this context. *TMK1* is also expressed in stomata and plays a significant role in abscisic acid (ABA) signaling (Yang et al. 2021), a key hormone regulating stomatal movement and water loss (Lim et al. 2015). Nonetheless, the role of TMKs in stomatal movement have never been systematically studied.

**Figure 7.**
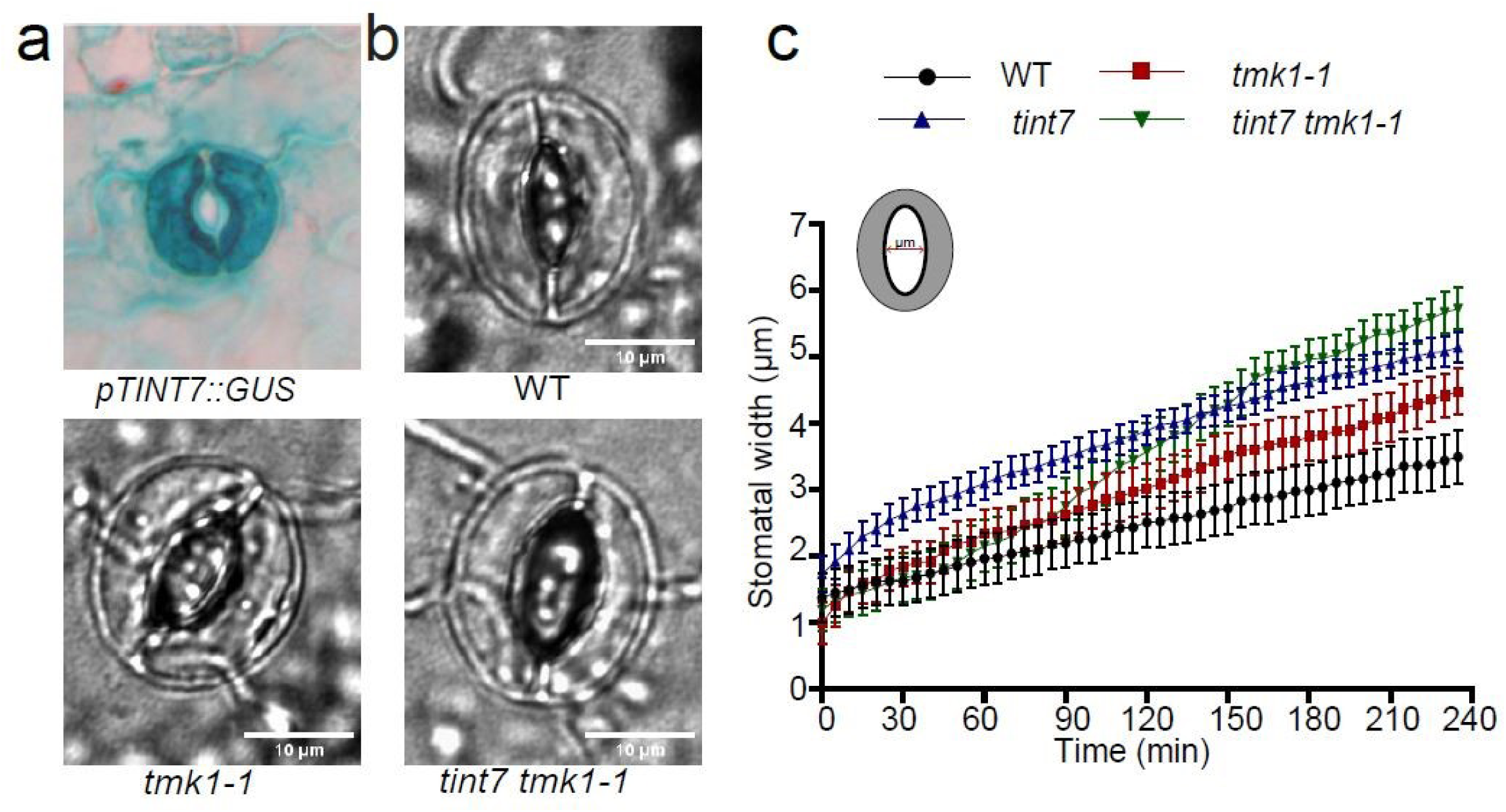
TINT7 function in regulating stomatal movement. (a) Representative image of *pTINT7::GUS* expression in stomata of the cotyledon of 5-day-old *A. thaliana* seedlings. (b) Representative images of stomatal opening after 4 hours of light in WT, *tmk1-1*, and *tint7 tmk1-1.* Scale bar, 10 µm. (c) Quantification of stomatal opening kinetics during light exposure. Stomatal width was measured every 5 minutes, as indicated in the scheme, and the average width with SEM is displayed in the graph. Stomatal opening in *tmk1-1* is greater than in WT, with *tint7* showing more opening than *tmk1-1*, *and tint7 tmk1-1* exhibiting the most pronounced opening. n >13 per genotype.

To explore a role of TINT7 in stomatal development or function, we first assessed the stomatal density in the *tint7* mutant and the *tint7 tmk1-1* double mutant. No significant differences in stomatal density were observed as compared to WT plants (Fig. S8a). Next, we evaluated the kinetics of stomatal movement (opening and closure) under different conditions. Following exposure to ABA, which promotes stomatal closure (Lim et al. 2015), the *tint7* mutant exhibited reduced closure compared to WT (Fig. S8b). Conversely, in response to light, the *tint7* mutant, similar to the *tmk1-1* single mutant, displayed greater stomatal opening compared to WT (Fig. 7b-c). This effect was more pronounced in the *tint7 tmk1-1* double mutant, which showed the highest rate and extent of stomatal opening (Fig. 7b-c).

Thus, the impaired stomatal movement phenotype in the *tint7* and *tint7 tmk1-1* mutants reveals a not yet well-characterized role of TMK signaling in this process.

## DISCUSSION

### TMK1 is part of large LRR-RLK interaction network

TMK1 is a notable receptor-like kinase involved in a broad range of essential plant functions. Its auxin sensing complex with ABP1 mediates the phosphorylation of about a thousand downstream targets (Friml et al. 2022; Kuhn et al. 2024). However, the precise mechanisms behind these phosphorylation events remain largely unknown and it is unclear whether this global phosphorylation response underlies all TMKs’ functions.

To gain deeper insights into TMKs’ roles and regulation, we investigated seven LRR-RLKs as potential TMK1 interactors (TINT), selected from the binary interaction network of LRR-RLKs (Smakowska-Luzan et al. 2018), and confirmed their association with TMK1 *in planta*. Here, we provide the initial characterization of this LRR-RLK TMK interacting network.

TINTs are evolutionary divergent within LRR-RLK family and classified into diverse subfamilies. For instance, TINT6 belongs to subfamily II, which consists mainly of co-receptors (Xi et al. 2019), some of which regulate meristem maintenance, anther development, and other functions. It is possible that TINT6 also acts as a co-receptor for TMK1 in various biological functions. In contrast, TINT7, being a larger receptor, likely has its own specific ligand, whose signaling converges with TMK-mediated auxin signaling. Notably, the alignment and analysis of the kinase domains revealed that all TINTs, except for TINT6 and TINT7, are likely pseudokinases. Due to their structural flexibility, pseudokinases can bring together multiple components of signaling networks or regulate active kinases allosterically, either enhancing or inhibiting their function (Zeqiraj and Van Aalten 2010; Sheetz and Lemmon 2022). This predominance of pseudokinases among TINT interactors raises intriguing questions about whether this pattern is coincidental or indicative of an evolutionary adaptation.

Regardless, the evolutionary, structural, and functional diversity of TINTs suggests that they were co-opted into the TMK pathway at several independent occasions during evolution and likely contribute to the diversity of functions and extensive regulation of TMK-based signaling.

### TINTs are part of TMK and CAMEL/CANAR regulatory network for auxin canalization

Auxin canalization is fundamental to plant development and survival in response to environmental challenges (Hajný et al. 2022). This unique property of auxin depends on the feedback regulation between its perception and transport, with ABP1/TMK auxin perception and PIN-mediated auxin transport serving as the respective principal components (Friml et al. 2022; Rodriguez et al. 2022; Wang et al. 2022). Nonetheless, the further mechanistic details remain poorly understood.

Vasculature regeneration around the wound and *de novo* vasculature formation from the local auxin source are preceded by formation of PIN1- and DR5-positive auxin channels; thus being classical manifestations of auxin canalization (Hajný et al. 2022). Among the TMK1 interactors, TINT3, TINT5, TINT6, and TINT7, based on the corresponding loss-of-function mutants, are required for these processes, similar to TMKs.

Two other LRR-RLKs, CAMEL and CANAR have been also identified as key components of auxin canalization (Hajný et al. 2020). Our findings revealed that TINT5 interacts not only with TMK1 but also with CAMEL, which, when considered alongside the very similar auxin canalization and vasculature regeneration defects observed in the *tmks*, *camel* and *tint5* mutants, highlight the potential importance of this interaction network in auxin canalization. Notably, the more pronounced canalization defects seen in the *tint5 tmk1-1* double mutant further emphasizes the need to investigate the functional implications of the TINT5 interaction with CAMEL and TMK1 in regulating auxin canalization.

Overall, the involvement of majority of TINTs along with TMKs in auxin canalization processes provides further genetic support for the interaction between TMKs and TINTs. Using TINT5 as an example, it also shows that the TINTs can link TMKs with other canalization components such as CAMEL and CANAR.

### TMK1 interactors suggest additional roles of TMK-based signaling

TMKs have been implicated in other processes beyond canalization, including apical hook opening, lateral root formation, leaf epidermal cell interdigitation, and others (Dai et al. 2013; Xu et al. 2014; Cao et al. 2019; Huang et al. 2019; Li et al. 2021; Marquès-Bueno et al. 2021; Rodriguez et al. 2022; Wang et al. 2022; Yu et al. 2023a).

Furthermore, other processes besides canalization involve auxin’s effect on PIN polarity. One of the best-characterized cases is the termination of shoot gravitropic bending, where, at later stages of the response, auxin accumulating at the lower side of the shoot leads to PIN3 repolarization, which is required for bending termination (Rakusová et al. 2016; Han et al. 2020). We found that TINT6 and TMK1 are involved in the gravitropic response of hypocotyls. The hyperbending observed in the *tint6* mutant suggests a disruption in the termination of bending. The rescue of this effect, or even reduced bending in the *tint6 tmk1-1* double mutant, implies a functional interaction between TINT6 and TMK1 in regulating the hypocotyl’s gravitropic response. These initial insights link auxin canalization and shoot bending termination genetically; however, the underlying mechanism and whether it involves PIN repolarization remain a topic for future investigation.

Characterization of *TINT7* revealed a connection to stomatal movement. Stomata and their regulated movement are key to regulating gas exchange and water balance, thus playing a vital role in photosynthesis and transpiration (Pillitteri and Dong 2013). We observed *TINT7* expression in stomata, and the *tint7* mutant showed reduced stomatal closure in response to ABA and enhanced opening in response to light, revealing TINT7’s role in stomatal movement. Additionally, the stomatal movement phenotype of *tmk1* and the more pronounced phenotype in the *tint7 tmk1-1* double mutant revealed a common function of TINT7 and TMK1 in this process. Further analysis will establish whether these mutants are more susceptible to water loss due to excessive stomatal opening and reduced responsiveness to ABA, a key hormone induced by osmotic stress during drought (Chua and Lau 2024). Thus, the TINT7 functional analysis provides the first insight into the role of TMK-mediated signaling in stomatal movement.

In summary, the functional analysis of various TINT proteins confirmed known and revealed new roles of TMKs in different developmental processes.

## ACKNOWLEDGEMENTS

We deeply appreciate M. Wrzaczek’s constructive input and insightful discussions, which significantly enriched this work. We thank L. Fiedler for helping with the heat map and for the discussions. We also thank the facilities at ISTA, the imaging and optics (IOF) and Lab Support (LSF) facilities for their service and assistance.

## AUTHOR CONTRIBUTIONs

A.M. took over the project, designed and executed the experiments, analyzed the data, and wrote the manuscript. L.R. contributed to the initial conceptualization and early stages of the project. E.M., M.Z., M.G., M.S., and E.Č. assisted with specific aspects of the methodology and data collection. J.F. conceptualized and supervised the study and contributed to the manuscript writing.

## METHODS

### Plant material, and growth conditions

The *Columbia-0* (*Col-0*) ecotype was the background of all the used lines of *Arabidopsis thaliana* (*A. thaliana*). Seeds were sterilized using chlorine gas overnight, followed by stratification for 48h in the dark at 4°C. Seedlings were grown on ½ MS medium plates (0.44% Murashige and Skoog basal salts, 1% sucrose, and 0.8% phyto-agar, pH 5.7) under long-day conditions (16 h light/ 8 h dark cycles, 22 ± 2°C) for 4 to 10 days, depending on specific assays, or in the soil under similar conditions. The T-DNA insertion line *tmk1-1* (SALK_016360) were previously reported(Cao et al. 2019). The other T-DNA insertion lines were obtained from NASC: *tint1* (SALKseq_053366), *tint2* (SALKseq_033657), *tint3* (SALKseq_086592), *tint4* (SALK_094070), *tint5* (SALKseq_131589), *tint5like* (SALK_078409C), *tint6* (SALK_123502C)*, tint7* (SALK_101617C).

The GUS lines of *pTINT1* through *TINT5* were obtained and previously reported (Wu et al. 2016). The line *pTMK1::GUS* was also previously reported (Friml et al. 2022).

### Plasmid construction and plant transformation

To generate the transgenic GUS lines of *TINT6* and *TINT7,* genomic fragments covering respectively 1000 bp and 1500 bp upstream from the start codon of *TINT6* and *TINT7* were amplified from the genomic DNA (gDNA) of *Col-0* and then inserted into the vector pGWB533. The resulting constructs were transformed into *Arabidopsis* plants by floral dipping in *Agrobacterium tumefaciens* cultures.

To generate the transgenic lines expressing HA-tagged TINT under ubiquitin promoter, the coding sequence (CDS) of each gene, without a stop codon, was amplified from *Col-0* complementary DNA (cDNA) through PCR with the corresponding primers attB1TINTCDS-Fw and attB2TINTnCDS-Rv (Supplementary table). This applies for *TINT1* through *TINT5*. By a BP reaction, the resulting sequences were inserted into pDONR221, which were then recombined with *pDONR P4-P1r pUBQ10* (Jaillais et al. 2011) and *pDONR P2R-P3 3xHA* into pB7m34GW vector by a MultiSite Gateway LR reaction. The resulting constructs were transformed into *Arabidopsis* plants by floral dipping in *Agrobacterium tumefaciens* cultures.

BiFC constructs were obtained by performing Gateway LR recombination of the gene entry clones with either pDEST–^GW^SCYNE for SCFP3A^N^ or pDEST–^GW^SCYCE for SCFP3A^C^ (Gehl et al. 2009).

*TINT6* and *TINT7* were amplified using primers with > 15bp overlapping sequences from *Col-0* cDNA, named TINT6/TINT7CDS-Fw and TINT6/TINT7CDS-Rv, designed for the Gibson Assembly method. The amplified sequences were assembled into the linearized *pDONR221* vector to generate the entry clone using the NEBuilder 409 HiFi DNA Assembly kit (E2621L); and then recombined with *pUBQ10* (Jaillais et al. 2011) and *pDONR P2R-P3 3xHA* into *pB7m34GW* vector by a MultiSite Gateway LR reaction.

To generate *pTINT5::TINT5-3HA*, *pTINT5-like::TINT5-like-3HA* constructs, we first amplified the promoter regions (1027 bp for *pTINT5* and 1292 bp for *pTINT5-like*) from *Col-0* gDNA with overlapping sequences. The amplified promoters were inserted into the linearized *pDONR P4-P1r* vector to generate the entry clone using the NEBuilder 409 HiFi DNA Assembly kit (E2621L). Similarly, *TINT5* and *TINT5-like* were amplified from *Col-0* cDNA with overlapping sequences, and then assembled into the linearized *pDONR221* vector. The entry clones of the promoter and the CDS regions were recombined by a Multisite Gateway LR reaction with *pDONR P2R-P3 3xHA* into *pB7m34GW* vector.

#### For FRET-FLIM

*p35S::CANAR-mCherry* and *p35S::CAMEL-mCherry* are previously reported (Hajný et al. 2020). To generate the construct p*35S::TINT5-GFP,* we used Gateway Cloning to recombine the entry vector of *TINT5* into *pB7FWG2*.

The lines *PIN2::PIN2-GFP*, *DR5rev::GFP* were described before (Benková et al. 2003; Friml et al. 2003). We generated *PIN2::PIN2-GFP* tint5/tin7 and *DR5rev::GFP tint5/tint7* by genetic crosses with *tint5/tint7.* The mutants *tint5* and *tint5-like* were crossed to generate the double mutant *tint5 tint5-like.* Equally, the double mutants of *tint5/tint6/tint6* were obtained by crosses with *tmk1-1*.

### Bioinformatics

The interaction network was built using Cytoscape (version 3.10.3, (Shannon et al. 2003)), based on the provided interaction data (Smakowska-Luzan et al. 2018). The data was imported into Cytoscape, where the network was visualized, analyzed, and further refined to depict potential interactions between the proteins of interest.

The protein sequence alignment was performed using ClustalW, a multiple sequence alignment tool (Madeira et al. 2024). Sequences were aligned according to the default settings of ClustalW.

The phylogenetic tree was constructed using a multiple sequence alignment, they were performed with the "build" function of ETE3 version 3.1.3 (Huerta-Cepas et al. 2016), as implemented on GenomeNet (https://www.genome.jp/tools/ete/). The tree was generated using FastTree with the slow nearest-neighbor interchange (NNI) method and MLACC=3 to make the maximum-likelihood NNIs more exhaustive (Price et al. 2010). The values at the nodes represent Shimodaira-Hasegawa (SH)-like local support.

The RNA-seq data was retrieved from the available online database on (https://plantrnadb.com/athrdb/) (Zhang et al. 2020). The expression data was processed and visualized in a heat map to display the median FPKM (fragments per kilobase of transcript per million mapped reads) expression of the genes of interest across different tissues. The heat map was generated using R.

### GUS staining

5-day-old seedlings were stained in 0.1 M sodium phosphate buffer (pH 7.0) containing 0.5 mM K_3_[Fe(CN)_6_], 0.5 mM K4[Fe(CN)6], and X-Gluc at 37°C for 1 h (*TMK1::GUS*), 1.5 h (*TINT2::GUS*, *TINT5::GUS*, *TINT6::GUS*), 2.5 h (*TINT1::GUS*, *TINT3::GUS*, *TINT4::GUS*) or 6 h (*TINT7::GUS*). Further, samples were incubated overnight in 80% (v/v) ethanol at room temperature. Tissue clearing was conducted as previously described (Malamy and Benfey 1997). DIC microscopy for analysis of the GUS staining assay was performed using an Olympus BX53 microscope equipped with 10x and 20x air objectives and a DP26 CCD camera.

### Vasculature regeneration after wounding in inflorescence stems

The regeneration experiments were performed as described previously (Mazur et al. 2016, 2020a, 2020b). Immature inflorescence stems (9–10 cm tall) of *Arabidopsis* were decapitated using a sharp razor blade. The apical floral parts (1–2 cm) were removed, and an artificial weight (a 2.5 g lead ball attached to a plastic tube) was applied to the stems. To prevent bending, decapitated stems were supported with a wooden stick. This setup allowed secondary tissue development in the basal parts of the immature stems (5-mm segments above the rosette) within six days of weight application.

For regeneration analysis, the inflorescence stems were precisely wounded with a razor blade about 5 mm from the rosette, in the transversal plane of the basal sectors with vascular cambium and secondary tissues. The plants remained covered with artificial weights throughout the experiment. Axillary buds above the rosette were not removed, serving as the source of endogenous auxin. Stem segments were cut at 0, 4, and 6 days after wounding using an automated vibratome (Leica VT1200 S), then sectioned at 70-µm-thick. Thee native sections were stained with 0.025% TBO aqueous solution, and regeneration was assessed in stems with fully developed, closed cambial rings and secondary tissues in the basal parts. The sections were observed under a bright-field microscope (Zeiss Axioscope.A1), and images of the vasculature were captured at 10× magnification using an Axiocam 506 camera.

The analysis of GFP reporter lines (*DR5rev::GFP* and *PIN1::PIN1-GFP*) was conducted 4 days after wounding using the Olympus FLUOVIEW FV1000 confocal laser-scanning microscope.

### GUS staining after wounding in inflorescence stems

For the observation of GUS activity after wounding, the same technique as used for regeneration analysis was applied. After 2, 4, and 6 days of wounding, stem segments were incubated in X-Gluc solution at 37°C. Afterward, they were fixed in 70% ethanol at room temperature. Samples with positive GUS reactions were sectioned at 70 µm using an automated vibratome. The native sections were then cleared in a solution containing 4% HCl and 20% methanol for 15 min at 65°C, followed by incubation in 7% NaOH and 70% ethanol for 15 min at room temperature. To rehydrate the samples, they were incubated in successive ethanol solutions (70%, 50%, 25%, and 10%) for 10 min each at room temperature, followed by a 10-min incubation in a solution of 25% glycerol and 5% ethanol. Finally, the seedlings were mounted in 50% glycerol and observed under a bright-field microscope. GUS activity images were captured at 10× magnification using a camera.

### Auxin-induced canalization in inflorescence stems (*de novo* vasculature formation)

The auxin canalization experiments were performed as described before (Mazur et al. 2020b), using *Arabidopsis* plants with young, 10-cm-tall inflorescence stems. Stems were wounded by making a transversal incision 3–4 mm above the rosette to disrupt the vascular cambium and secondary tissues, thereby interrupting the polar basipetal auxin transport. Then an exogenous auxin application of 10 µM IAA (Sigma-Aldrich, 15148-2G) in a droplet of lanolin paste was applied below the cut, which was replaced every two days to maintain a consistent auxin presence. Samples were collected, and longitudinal stem sections were manually prepared using a NIKON SMZ1500 stereomicroscope. The sections were stained with 0.05% TBO and mounted in 50% glycerol aqueous solution. Images were captured using an Olympus BX43 microscope with an Olympus SC30 Camera.

The analysis of GFP reporter lines (*DR5rev::GFP* and *PIN1::PIN1-GFP*) was conducted 4 days after local applications using the Olympus FLUOVIEW FV1000 confocal laser-scanning microscope.

### Cotyledon vasculature analysis

To observe vein patterns in cotyledons, shoot tissues, including cotyledons from 10-day-old seedlings of each genotype, were harvested and cleared in 70% ethanol for 3 days, with the solution being changed intermittently. The samples were then incubated cleared in a solution containing 4% HCl and 20% methanol for 15 min at 65°C, followed by incubation in 7% NaOH and 70% ethanol for 15 min at room temperature. To rehydrate the samples, they were incubated in successive ethanol solutions (70%, 50%, 25%, and 10%) for 10 min each at room temperature, followed by a 10-min incubation in a solution of 25% glycerol and 5% ethanol. Finally, the seedlings were mounted in 50% glycerol and observed under a stereomicroscope (Olympus SZX16 Stereo).

Vein patterns in the cotyledons were evaluated and categorized as either defective or non-defective. Cotyledons with exactly four closed loops were considered non-defective. Additionally, cotyledons were deemed non-defective if any of the two lower loops were open from the bottom. Conversely, cotyledons exhibiting more or fewer than four loops, any extra branches, or open loops, were classified as defective. The number of analyzed cotyledons was between 80 and 100 per genotype.

### Tobacco (Nicotiana bethamiana) infiltration

Constructs were transformed into *Agrobacterium tumefaciens* GV3101 and grown on LB plates at 28°C for 48 hours. Subsequently, liquid LB suspensions carrying the desired plasmids were grown overnight at 28-30 °C, after which young tobacco leaves were infiltrated or co-infiltrated. Additionally to an overnight in the dark, the plants were grown under normal conditions for 36 hours. The infiltatred leaves were then either harvested for co-immunoprecipitation assay or used for bimolecular fluorescence complementation (BiFC) imaging.

### Bimolecular fluorescence complementation (BiFC) assay in *Nicotiana benthamiana*

BiFC assays in *Nicotiana benthamiana* were performed by co-infiltrating the leaves with *p35S::TMK1-SCFP3A^N^* and *p35S::TINT-SCFP3A^C^*, *p35S::TMK1-SCFP3A^C^*, *or p35S::TMK2-SCFP3A^C^*. Imaging was then performed two days after using Zeiss LSM800 microscope.

### Co-immunoprecipitation (Co-IP) assay

To immunoprecipitate HA-tagged TINT proteins, either 5-day old *Arabidopsis* seedlings from the corresponding transgenic line (expressing HA-tagged TINT under ubiquitin promoter) or leaves from infiltrated or co-infiltrated *Nicotiana benthamiana* (see above) were harvested, ground into powder using liquid nitrogen, and homogenized in the lysis buffer (150 mM NaCl, 1% Triton X-100, 50 mM Tris HCl pH 8, Complete protease cocktail and PhosStop phosphatase inhibitor cocktail (Roche)). The total protein extract is obtained after 30 minutes centrifugation at 14000 g, 4°C. Imunoprecipitation was then performed following the manufacturer’s instructions (Miltenyi Biotec) using microMACS beads coupled to a monoclonal anti-HA antibody. The immunoprecipitated proteins antibody were separated by SDS-PAGE using 10% Mini-PROTEAN®TGX™ Precast Protein Gels (Bio-RAD), transferred to a PVDF membrane, and analyzed by immunoblotting. TINT-HA was detected using anti-HA HRP-conjugated, High Affinity (3F10) antibody (Roche), while co-immunoprecipitated TMK1-FLAG from *Nicotiana benthamiana* was detected with anti-FLAG® M2-Peroxidase (HRP) antibody (Sigma). For *Arabidopsis* samples, endogenous TMK1 was detected using anti-TMK1 antibody (Nordic Biosite), which was incubated overnight at 4°C, followed by incubation with anti-rabbit HRP antibody at room temperature for 2 hours. Signal detection was performed using the SuperSignal West Femto Maximum Sensitivity Substrate Detection System (Thermo Scientific), and images were captured with an Amersham 600RGB 604 image analyzer (GE Healthcare).

### FRET-FLIM in *Arabidopsis* root suspension culture protoplasts

Protoplasts from *Arabidopsis* root cell suspension cultures were isolated using a PEG-mediated transformation method. Three-day-old cells were collected by centrifugation into a 50 ml Falcon tube and incubated for 3-4 hours in dark in a petri dish containing an enzyme solution. This solution included 1% of Cellulase RS, 0.2% macroenzyme dissolved in the B5-0.34M glucose-mannitol (GM) solution. Following a 5-minute centrifugation and two washes with the GM solution, the cells were transferred to a 0.28M sucrose solution in a 15ml Falcon tube and floated by spinning at 800 rpm for 7 minutes. The floating cells were then collected into a microcentrifuge tube and diluted to a final concentration of 4×10^^6^ cells/mL using the GM solution.

For transfection, 50 μL of protoplasts were mixed with 10 μg of high-purity plasmid DNA corresponding to the appropriate vectors and 150 μL of PEG buffer. The mixture was incubated in the dark at room temperature for 60 minutes, followed by a single wash with Ca(NO3)_2_ solution. The transfected cells were then resuspended in 500 μL of GM solution, transferred to a 24-well plate, and incubated overnight at room temperature in the dark before imaging.

Imaging of FRET-FLIM experiments was performed the next day using a Leica SP8 equipped with a FALCON FLIM detector. Fluorescence lifetime measurements were acquired using LAS X software (FLIM).

### Immunolocalization

Whole-mount *in situ* immunolocalization was performed on 4d old seedlings of *Arabidopsis* following the published protocol (Sauer and Friml 2010).The anti-HA antibody (Thermo Fisher, Monoclonal Antibody (2-2.2.14)) was applied at a 1:500 dilution, and the secondary Cy3 Anti-Mouse IgG (Sigma-Aldrich, Sheep Anti-Mouse IgG) was used at a 1:600 dilution. Imaging of the immuno-stained roots was performed using Zeiss LSM800 microscope (Cy3: excitation at 548 nm and emission at 561 nm).

### Hypocotyl gravitropism assays

Plates containing *Arabidopsis* seeds were placed vertically in the growth room under light for 6 hours to induce germination. The plates were then covered with aluminum foil and placed in a box to maintain darkness for an additional 4 days. Following this, the seedlings were subjected to a 90-degree rotation to provide gravity stimulation. After 24 hours, the plates were scanned using an EPSON V700 scanner, and the bending angle of the seedlings was measured using ImageJ.

### Stomata patterning

Stomatal patterning was analyzed using cotyledons from 10-day-old seedlings. The cotyledons were placed on a slide with water and covered with a cover slip, then imaged using an Olympus BX53 microscope equipped with 10x and 20x air objectives and a DP26 CCD camera. The number of stomata was quantified within a defined area of consistent size across all samples using ImageJ software.

### Stomatal movement assays

To prepare samples for stomatal movement assays, healthy 7^th^ or 8^th^ rosette leaves were selected from 4-week-old *Arabidopsis* plants. Leaves were carefully excised to prevent tissue damage, cut into small pieces, and the lower epidermal peel was prepared. The isolated epidermal peels were placed in a 24-well plate, with double-sided tape in each well and a bottom cover slip. Buffer solutions were added to the wells depending on the assay. For the light-induced opening assay, the buffer contained 10 mM MES-KOH (pH 6.15) and 10 mM KCl to induce stomatal closure overnight. For the ABA treatment assay, the buffer included10 mM MES-KOH (pH 6.15), 30 mM KCL and 50 μM CaCl_2_. The plate was incubated overnight at 22°C under plant growth conditions (150 µmol/m²/s light intensity, 16-hour light/8-hour dark cycle) to stabilize the samples.

The following morning, 3 hours into the circadian light cycle, the epidermal peels were transferred to fresh buffer solutions corresponding to their respective treatments. For the ABA assay, 10 μM ABA was added to induce stomatal closure. For the light-induced opening assay, the peels were exposed to direct light to trigger stomatal opening. Stomatal movements were observed using a Nikon Ti2E inverted microscope with a 40× objective, capturing images every 5 minutes over a 4-hour period. Stomatal widths were measured throughout the experiment using ImageJ image analysis software.

## REFERENCES

Balla J, Kalousek P, Reinöhl V, Friml J, and Procházka S. Competitive canalization of PIN-dependent auxin flow from axillary buds controls pea bud outgrowth. Plant J. 2011:65(4):571–577. 10.1111/j.1365-313X.2010.04443.x

Benková E, Michniewicz M, Sauer M, Teichmann T, Seifertová D, Jürgens G, and Friml J. Local, Efflux-Dependent Auxin Gradients as a Common Module for Plant Organ Formation. Cell. 2003:115(5):591–602. 10.1016/s0092-8674(03)00924-3

Cao M, Chen R, Li P, Yu Y, Zheng R, Ge D, Zheng W, Wang X, Gu Y, Gelová Z, et al. TMK1-mediated auxin signalling regulates differential growth of the apical hook. Nature. 2019:568(7751):240–243. 10.1038/s41586-019-1069-7

Chang C, Schaller GE, Patterson SE, Kwok SF, Meyerowitz EM, and Bleecker AB. The TMK1 gene from Arabidopsis codes for a protein with structural and biochemical characteristics of a receptor protein kinase. Plant Cell. 1992:4(10):1263–1271. 10.1105/tpc.4.10.1263

Chen X, Wang T, Rehman AU, Wang Y, Qi J, Li Z, Song C, Wang B, Yang S, and Gong Z. Arabidopsis U-box E3 ubiquitin ligase PUB11 negatively regulates drought tolerance by degrading the receptor-like protein kinases LRR1 and KIN7. JIPB. 2021:63(3):494–509. 10.1111/jipb.13058

Chinchilla D, Zipfel C, Robatzek S, Kemmerling B, Nürnberger T, Jones JDG, Felix G, and Boller T. A flagellin-induced complex of the receptor FLS2 and BAK1 initiates plant defence. Nature. 2007:448(7152):497–500. 10.1038/nature05999

Chua LC and Lau OS. Stomatal development in the changing climate. Development. 2024:151(20):dev202681. 10.1242/dev.202681

Dai N, Wang W, Patterson SE, and Bleecker AB. The TMK Subfamily of Receptor-Like Kinases in Arabidopsis Display an Essential Role in Growth and a Reduced Sensitivity to Auxin. PLoS ONE. 2013:8(4). 10.1371/journal.pone.0060990

Escocard De Azevedo Manhães AM, Ortiz-Morea FA, He P, and Shan L. Plant plasma membrane-resident receptors: Surveillance for infections and coordination for growth and development. JIPB. 2021:63(1):79–101. 10.1111/jipb.13051

Friml J. Fourteen Stations of Auxin. Cold Spring Harb Perspect Biol. 2022:14(5). 10.1101/cshperspect.a039859

Friml J, Gallei M, Gelová Z, Johnson A, Mazur E, Monzer A, Rodriguez L, Roosjen M, Verstraeten I, Živanović BD, et al. ABP1–TMK auxin perception for global phosphorylation and auxin canalization. Nature. 2022:609(7927):575–581. 10.1038/s41586-022-05187-x

Friml J, Vieten A, Sauer M, Weijers D, Schwarz H, Hamann T, Offringa R, and Jürgens G. Efflux-dependent auxin gradients establish the apical–basal axis of Arabidopsis. Nature. 2003:426(6963):147–153. 10.1038/nature02085

Gallei M, Luschnig C, and Friml J. Auxin signalling in growth: Schrödinger’s cat out of the bag. Curr Opin Plant Biol. 2020:53:43–49. 10.1016/j.pbi.2019.10.003

Gehl C, Waadt R, Kudla J, Mendel R-R, and Hänsch R. New GATEWAY vectors for High Throughput Analyses of Protein–Protein Interactions by Bimolecular Fluorescence Complementation. Molecular Plant. 2009:2(5):1051–1058. 10.1093/mp/ssp040

Gu B, Dong H, Smith C, Cui G, Li Y, and Bevan MW. Modulation of receptor-like transmembrane kinase 1 nuclear localization by DA1 peptidases in Arabidopsis. Proc Natl Acad Sci USA. 2022:119(40). 10.1073/pnas.2205757119

Hajný J, Prát T, Rydza N, Rodriguez L, Tan S, Verstraeten I, Domjan D, Mazur E, Smakowska-Luzan E, Smet W, et al. Receptor kinase module targets PIN-dependent auxin transport during canalization. Science. 2020:370(6516):550–557. 10.1126/science.aba3178

Hajný J, Tan S, and Friml J. Auxin canalization: From speculative models toward molecular players. Curr Opin Plant Biol. 2022:65. 10.1016/j.pbi.2022.102174

Han H, Rakusová H, Verstraeten I, Zhang Y, and Friml J. SCF^TIR1/AFB^ Auxin Signaling for Bending Termination during Shoot Gravitropism. Plant Physiol. 2020:183(1):37–40. 10.1104/pp.20.00212

Hohmann U, Lau K, and Hothorn M. The Structural Basis of Ligand Perception and Signal Activation by Receptor Kinases. Annu Rev Plant Biol. 2017:68(1):109–137. 10.1146/annurev-arplant-042916-040957

Hu C, Zhu Y, Cui Y, Cheng K, Liang W, Wei Z, Zhu M, Yin H, Zeng L, Xiao Y, et al. A group of receptor kinases are essential for CLAVATA signalling to maintain stem cell homeostasis. Nature Plants. 2018:4(4):205–211. 10.1038/s41477-018-0123-z

Huang R, Zheng R, He J, Zhou Z, Wang J, Xiong Y, and Xu T. Noncanonical auxin signaling regulates cell division pattern during lateral root development. Proc Natl Acad Sci USA. 2019:116(42):21285–21290. 10.1073/pnas.1910916116

Huerta-Cepas J, Serra F, and Bork P. ETE 3: Reconstruction, Analysis, and Visualization of Phylogenomic Data. Mol Biol Evol. 2016:33(6):1635–1638. 10.1093/molbev/msw046

Jaillais Y, Hothorn M, Belkhadir Y, Dabi T, Nimchuk ZL, Meyerowitz EM, and Chory J. Tyrosine phosphorylation controls brassinosteroid receptor activation by triggering membrane release of its kinase inhibitor. Genes Dev. 2011:25(3):232–237. 10.1101/gad.2001911

Jiang J, Zhang C, and Wang X. Ligand Perception, Activation, and Early Signaling of Plant Steroid Receptor Brassinosteroid Insensitive 1. JIPB. 2013:55(12):1198–1211. 10.1111/jipb.12081

Jose J, Ghantasala S, and Roy Choudhury S. Arabidopsis Transmembrane Receptor-Like Kinases (RLKs): A Bridge between Extracellular Signal and Intracellular Regulatory Machinery. IJMS. 2020:21(11):4000. 10.3390/ijms21114000

Kuhn A, Roosjen M, Mutte S, Dubey SM, Carrasco VPC, Boeren S, Monzer A, Koehorst J, Kohchi T, Nishihama R, et al. RAF-like protein kinases mediate a deeply conserved, rapid auxin response. Cell. 2024:187(1):130–148. 10.1016/j.cell.2023.11.021

Li B, Zhou Q, Cai L, Li L, Xie C, Li D, Zhu F, Li X, Zhao X, Liu X, et al. TMK4-mediated FIP37 phosphorylation regulates auxin-triggered N-methyladenosine modification of auxin biosynthetic genes in Arabidopsis. Cell Reports. 2024:43(8):114597. 10.1016/j.celrep.2024.114597

Li L, Verstraeten I, Roosjen M, Takahashi K, Rodriguez L, Merrin J, Chen J, Shabala L, Smet W, Ren H, et al. Cell surface and intracellular auxin signalling for H+ fluxes in root growth. Nature. 2021:599(7884):273–277. 10.1038/s41586-021-04037-6

Lim C, Baek W, Jung J, Kim J-H, and Lee S. Function of ABA in Stomatal Defense against Biotic and Drought Stresses. IJMS. 2015:16(7):15251–15270. 10.3390/ijms160715251

Liu J, Li W, Wu G, and Ali K. An update on evolutionary, structural, and functional studies of receptor-like kinases in plants. Front Plant Sci. 2024:15:1305599. 10.3389/fpls.2024.1305599

Liu P-L, Du L, Huang Y, Gao S-M, and Yu M. Origin and diversification of leucine-rich repeat receptor-like protein kinase (LRR-RLK) genes in plants. BMC Evol Biol. 2017:17(1):47. 10.1186/s12862-017-0891-5

Luschnig C and Friml J. Over 25 years of decrypting PIN-mediated plant development. Nat Commun. 2024:15(1):9904. 10.1038/s41467-024-54240-y

Madeira F, Madhusoodanan N, Lee J, Eusebi A, Niewielska A, Tivey ARN, Lopez R, and Butcher S. The EMBL-EBI Job Dispatcher sequence analysis tools framework in 2024. Nucleic Acids Research. 2024:52(W1):W521–W525. 10.1093/nar/gkae241

Malamy JE and Benfey PN. Organization and cell differentiation in lateral roots of *Arabidopsis thaliana*. Development. 1997:124(1):33–44. 10.1242/dev.124.1.33

Marquès-Bueno MM, Armengot L, Noack LC, Bareille J, Rodriguez L, Platre MP, Bayle V, Liu M, Opdenacker D, Vanneste S, et al. Auxin-Regulated Reversible Inhibition of TMK1 Signaling by MAKR2 Modulates the Dynamics of Root Gravitropism. Curr Biol. 2021:31(1):228–237.e10. 10.1016/j.cub.2020.10.011

Mazur E, Benková E, and Friml J. Vascular cambium regeneration and vessel formation in wounded inflorescence stems of Arabidopsis. Sci Rep. 2016:6. 10.1038/srep33754

Mazur E, Gallei M, Adamowski M, Han H, Robert HS, and Friml J. Clathrin-mediated trafficking and PIN trafficking are required for auxin canalization and vascular tissue formation in Arabidopsis. Plant Science. 2020a:293:110414. 10.1016/j.plantsci.2020.110414

Mazur E, Kulik I, Hajný J, and Friml J. Auxin canalization and vascular tissue formation by TIR1/AFB-mediated auxin signaling in Arabidopsis. New Phytologist. 2020b:226(5):1375–1383. 10.1111/nph.16446

Pillitteri LJ and Dong J. Stomatal Development in Arabidopsis. The Arabidopsis Book. 2013:11:e0162. 10.1199/tab.0162

Price MN, Dehal PS, and Arkin AP. FastTree 2 – Approximately Maximum-Likelihood Trees for Large Alignments. PLoS ONE. 2010:5(3):e9490. 10.1371/journal.pone.0009490

Rakusová H, Abbas M, Han H, Song S, Robert HS, and Friml J. Termination of Shoot Gravitropic Responses by Auxin Feedback on PIN3 Polarity. Current Biology. 2016:26(22):3026–3032. 10.1016/j.cub.2016.08.067

Rodriguez L, Fiedler L, Zou M, Giannini C, Monzer A, Gelová Z, Verstraeten I, Hajný J, Tan S, Hoermayer L, et al. Cell surface auxin signalling directly targets PIN-mediated auxin fluxes for adaptive plant development. 2022. 10.1101/2022.11.30.518503

Sachs T. The Control of the Patterned Differentiation of Vascular Tissues.. In. Advances in Botanical Research. (Elsevier), pp. 151–262. 10.1016/S0065-2296(08)60351-1

Salehin M, Bagchi R, and Estelle M. SCF^TIR1/AFB^ -Based Auxin Perception: Mechanism and Role in Plant Growth and Development. Plant Cell. 2015:27(1):9–19. 10.1105/tpc.114.133744

Sauer M and Friml J. Immunolocalization of Proteins in Plants.. In. Plant Developmental Biology, L Hennig and C Köhler, eds, Methods in Molecular Biology. (Humana Press: Totowa, NJ), pp. 253–263. 10.1007/978-1-60761-765-5_17

Scarpella E, Marcos D, Friml J, and Berleth T. Control of leaf vascular patterning by polar auxin transport. Genes Dev. 2006:20(8):1015–1027. 10.1101/gad.1402406

Shannon P, Markiel A, Ozier O, Baliga NS, Wang JT, Ramage D, Amin N, Schwikowski B, and Ideker T. Cytoscape: A Software Environment for Integrated Models of Biomolecular Interaction Networks. Genome Res. 2003:13(11):2498–2504. 10.1101/gr.1239303

Sheetz JB and Lemmon MA. Looking lively: emerging principles of pseudokinase signaling. Trends in Biochemical Sciences. 2022:47(10):875– 891. 10.1016/j.tibs.2022.04.011

Shiu S-H and Bleecker AB. Plant Receptor-Like Kinase Gene Family: Diversity, Function, and Signaling. Sci STKE. 2001:2001(113). 10.1126/stke.2001.113.re22

Smakowska-Luzan E, Mott GA, Parys K, Stegmann M, Howton TC, Layeghifard M, Neuhold J, Lehner A, Kong J, Grünwald K, et al. An extracellular network of Arabidopsis leucine-rich repeat receptor kinases. Nature. 2018:553(7688):342–346. 10.1038/nature25184

Takeuchi H and Higashiyama T. Tip-localized receptors control pollen tube growth and LURE sensing in Arabidopsis. Nature. 2016:531(7593):245–248. 10.1038/nature17413

Wang J, Chang M, Huang R, Gallei M, Friml J, Yu Y, Wen M, Yang Z, and Xu T. Self-regulation of PIN1-driven auxin transport by cell surface-based auxin signaling in Arabidopsis. 2022. 10.1101/2022.11.30.518523

Wang JL, Wang M, Zhang L, Li YX, Li JJ, Li YY, Pu ZX, Li DY, Liu XN, Guo W, et al. WAV E3 ubiquitin ligases mediate degradation of IAA32/34 in the TMK1-mediated auxin signaling pathway during apical hook development. Proc Natl Acad Sci USA. 2024:121(17). 10.1073/pnas.2314353121

Wang Q, Qin G, Cao M, Chen R, He Y, Yang L, Zeng Z, Yu Y, Gu Y, Xing W, et al. A phosphorylation-based switch controls TAA1-mediated auxin biosynthesis in plants. Nat Commun. 2020:11(1). 10.1038/s41467-020-14395-w

Wu Y, Xun Q, Guo Y, Zhang J, Cheng K, Shi T, He K, Hou S, Gou X, and Li J. Genome-Wide Expression Pattern Analyses of the Arabidopsis Leucine-Rich Repeat Receptor-Like Kinases. Molecular Plant. 2016:9(2):289–300. 10.1016/j.molp.2015.12.011

Xi L, Wu XN, Gilbert M, and Schulze WX. Classification and Interactions of LRR Receptors and Co-receptors Within the Arabidopsis Plasma Membrane – An Overview. Front Plant Sci. 2019:10:472. 10.3389/fpls.2019.00472

Xu T, Dai N, Chen J, Nagawa S, Cao M, Li H, Zhou Z, Chen X, De Rycke R, Rakusová H, et al. Cell Surface ABP1-TMK Auxin-Sensing Complex Activates ROP GTPase Signaling. Science. 2014:343(6174):1025–1028. 10.1126/science.1245125

Yang J, He H, He Y, Zheng Q, Li Q, Feng X, Wang P, Qin G, Gu Y, Wu P, et al. TMK1-based auxin signaling regulates abscisic acid responses via phosphorylating ABI1/2 in Arabidopsis. Proc Natl Acad Sci USA. 2021:118(24). 10.1073/pnas.2102544118

Yu Y, Tang W, Lin W, Li W, Zhou X, Li Y, Chen R, Zheng R, Qin G, Cao W, et al. ABLs and TMKs are co-receptors for extracellular auxin. Cell. 2023a:186(25):5457–5471.e17. 10.1016/j.cell.2023.10.017

Yu Z, Ma J, Zhang M, Li X, Sun Y, Zhang M, and Ding Z. Auxin promotes hypocotyl elongation by enhancing BZR1 nuclear accumulation in *Arabidopsis*. Sci Adv. 2023b:9(1):eade2493. 10.1126/sciadv.ade2493

Zeqiraj E and Van Aalten DM. Pseudokinases-remnants of evolution or key allosteric regulators? Current Opinion in Structural Biology. 2010:20(6):772–781. 10.1016/j.sbi.2010.10.001

Zhang H, Zhang F, Yu Y, Feng L, Jia J, Liu B, Li B, Guo H, and Zhai J. A Comprehensive Online Database for Exploring ∼20,000 Public Arabidopsis RNA-Seq Libraries. Molecular Plant. 2020:13(9):1231–1233. 10.1016/j.molp.2020.08.001

Zhu Q, Feng Y, Xue J, Chen P, Zhang A, and Yu Y. Advances in Receptor-like Protein Kinases in Balancing Plant Growth and Stress Responses. Plants. 2023:12(3):427. 10.3390/plants12030427

