## Supplementary figures and table for "TMK interacting network of receptor like kinases for auxin canalization and beyond"

a

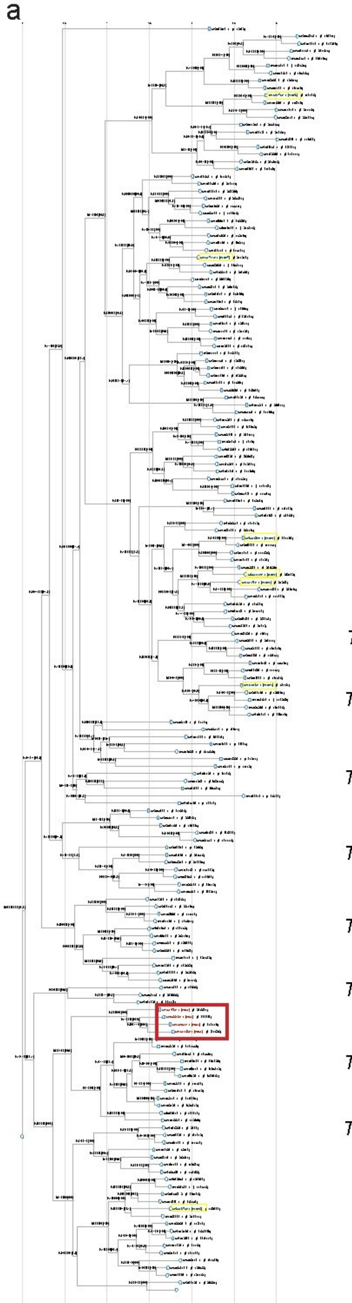

b

| Name | ATG | LRR-RLK sub-family | Size (aa) |
|---|---|---|---|
| *TMK1* | AT1G66150 | IX | 942 |
| *TINT1 (LRR1)* | AT5G16590 | III | 625 |
| *TINT2* | AT4G23740 | III | 638 |
| *TINT3 (PRK7)* | AT4G31250 | III | 676 |
| *TINT4* | AT5G24100 | III | 614 |
| *TINT5* | AT2G02780 | VI | 742 |
| *TINT6 (CIK4)* | AT5G45780 | II | 614 |
| *TINT7* | AT1G17230 | XI | 1101 |

c

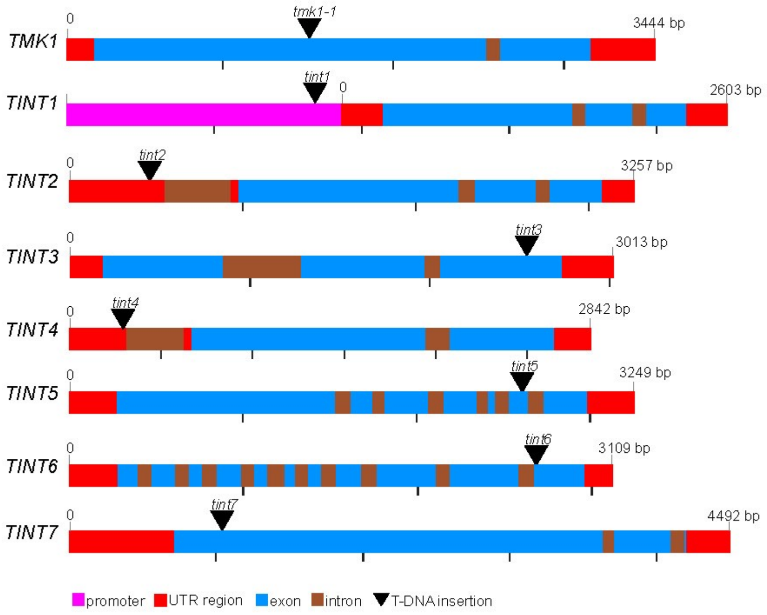

**Figure S1- Phylogenetic analysis and characterization of TINT in *Arabidopsis***

(a) Phylogenetic tree of LRR-RLKs in *Arabidopsis*, with the TMK family highlighted in red boxes and the TINT family highlighted in yellow boxes. The tree was constructed using FastTree with maximum-likelihood analysis. Node values represent Shimodaira-Hasegawa (SH)-like local support, with higher values reflecting stronger confidence in the tree structure.

(b) Table summarizing the TINT proteins, including their gene identifiers (ATG), retrieved from the *Arabidopsis* Information Resource (TAIR), their LRR-RLK subfamily, and protein size (in amino acids).

(c) Schematic representation of the genomic DNA (gDNA) regions of *TMK1* and *TINT,* with the transcriptional start site marked as zero. The black triangle indicates the site of the T-DNA insertion in the mutants.

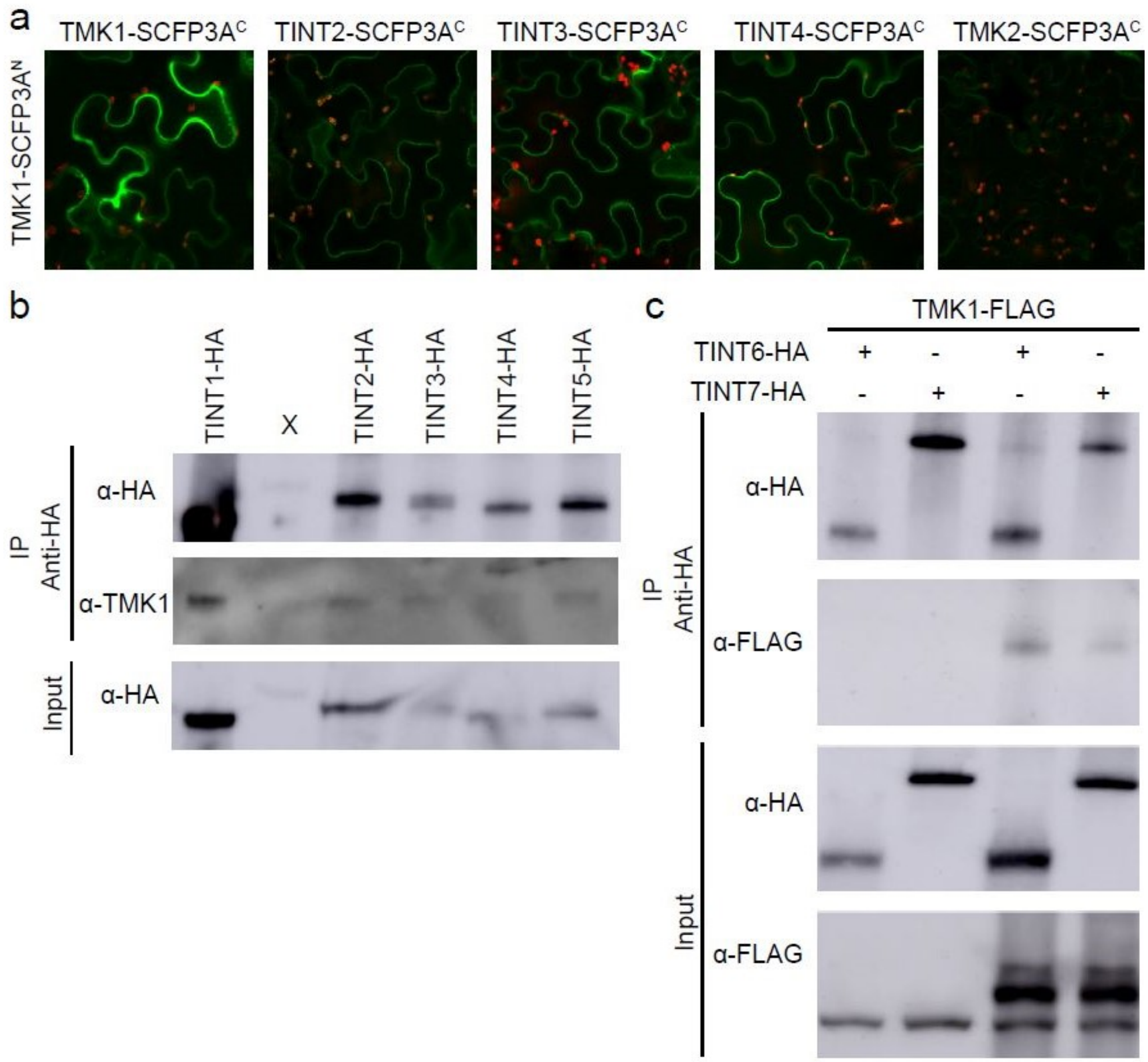

**Figure S2- Confirmation for the interaction of TINTs with TMK1**

(a) BiFC assay performed in *Nicotiana benthamiana* leaves co-expressing constructs for split the super cyan fluorescent protein (SCFP3A) by fusing the N-terminus of SCFP3A to TMK1 and the C-terminus to the protein of interest. Protein-protein interaction was assessed by fluorescence complementation. The complementation signal was observed with TINT2, TINT3, and TINT4. TMK1-SCFP3A$^N$ served as a positive control, while TMK2-SCFP3A$^N$ was used as a negative control.

(b) Co-immunoprecipitation assay from *Arabidopsi*s seedlings expressing the corresponding TINT with HA tag under the ubiquitin promoter (*pUBQ10::TINTn-3HA*). TINT proteins were immunoprecipitated using HA antibody beads, and an antibody against TMK1 was used to assess co-immunoprecipitation. While the interaction is observed, the bands are weak due to the use of an endogenous antibody.

(c) Co-immunoprecipitation assay from *Nicotiana benthamiana* leaves co-expressing *p35S::TMK1-FLAG* with *pUBQ10::TINT6/TINT7-3HA*. TINT proteins were immunoprecipitated using HA antibody beads, and an antibody against FLAG was used to assess co-immunoprecipitation. The interaction between TINT6/TINT7 and TMK1 was observed.

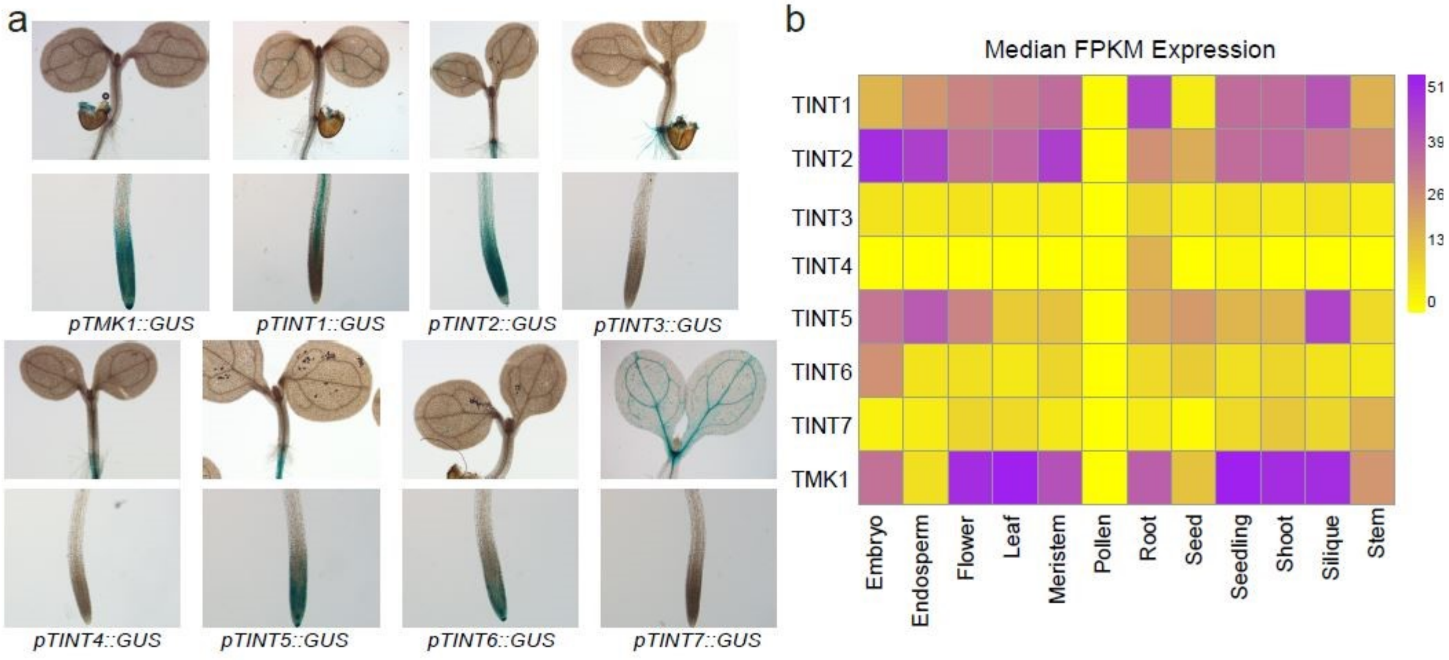

**Figure S3-Expression pattern of TINTs**

(a) Representative images of the GUS staining of 5-day-old *Arabidopsis* seedlings showing different expression of TINTs across the root, shoot, and cotyledons.
(b) Heat map of the expression profiles of *TMK1* and *TINT* genes, showing the median FPKM expression values across different plant tissues. Expression levels range from 0 (no detectable expression) to 51 (high expression).

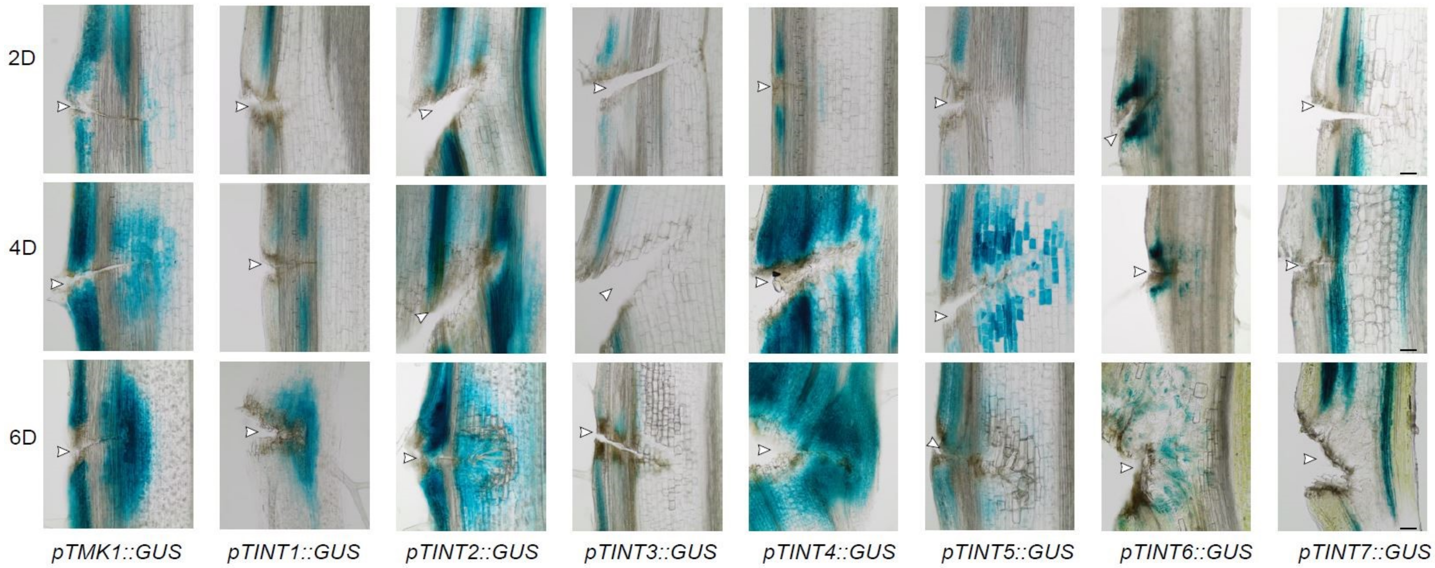

**Figure S4- TINT expression during vasculature regeneration**

Representative images of the GUS staining of inflorescence stems during vasculature regeneration at 2, 4 and 6 days after wounding (DAW) in GUS lines. White arrowheads indicate the wounding site. Scale bar, 100 µm.

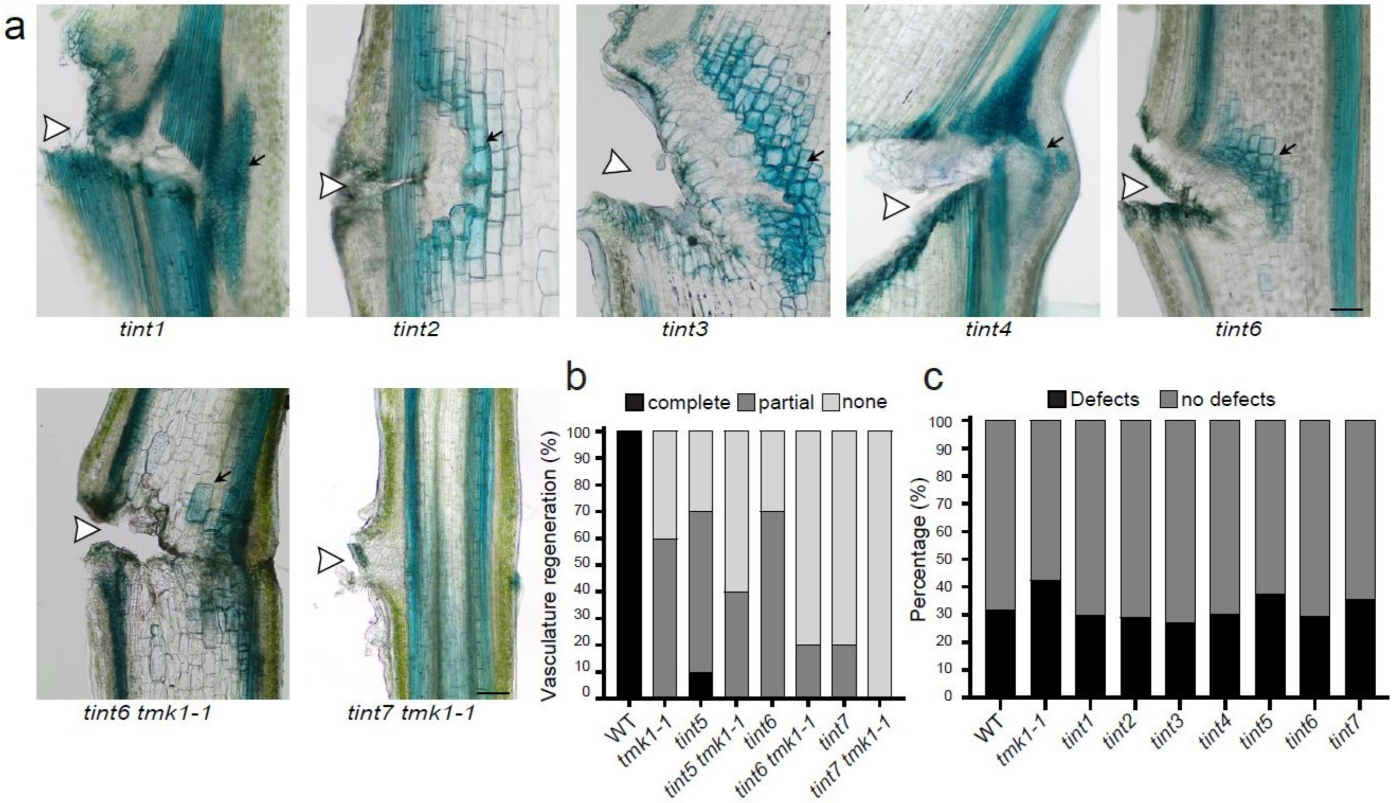

**Figure S5- TINT and TMK1 in vasculature regeneration after wounding**

(a) Representative images of the TBO staining at 6 DAW showing vasculature regeneration in the inflorescence stems. Stems of *tint1*, *tint2*, and *tint4* regenerated fully around the wound, while *tint3* and *tint6* showed partial regeneration. The *tint6 tmk1-1* double mutant developed single vessels, and the *tint7 tmk1-1* mutant showed no vasculature regeneration. White arrowheads indicate the wounding site. Black arrows indicate the regenerated vasculature. Scale bar, 100 µm.

(b) Quantification of the vasculature regeneration in the single and double mutants, categorized as complete (fully formed vasculature), partial (limited and partially developed vasculature), or none (no vessel formed around the wound) at 6 DAW. n=10 plants per genotype.

(c) Quantification of the cotyledon vasculature phenotype, categorized as no defects (4 loops, all closed), or defects (fewer or more than 4 loops, or open loops except for the bottom loops open at the bottom). All mutants exhibited normal cotyledon vasculature development, similar to WT. n >100 per genotype.

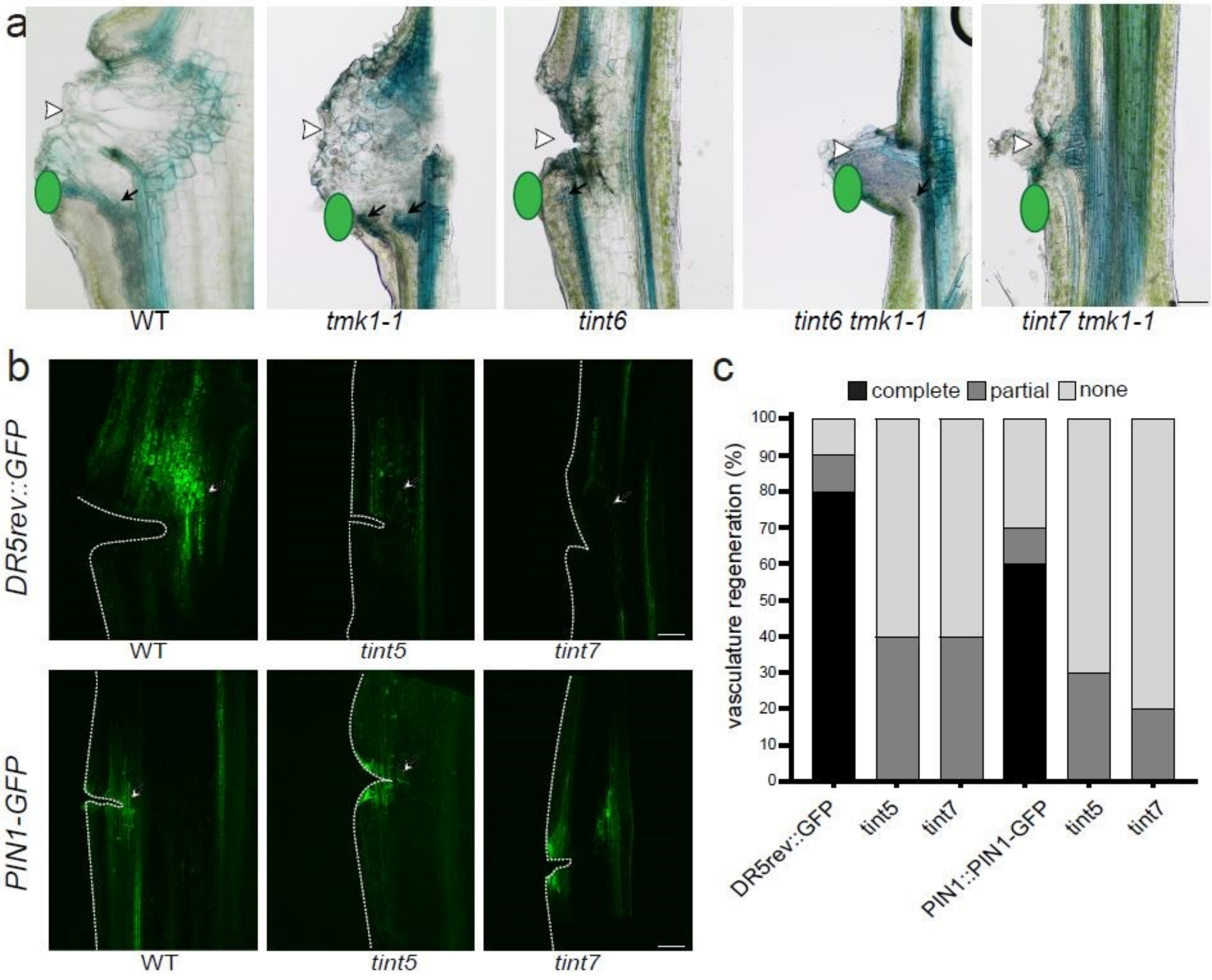

**Figure S6- TINT and TMK1 in auxin channel formation**

(a) Representative images of the TBO staining of *de novo* vasculature formation and canalization at 6 days after IAA application (DAA) (green ovals) following wounding. WT vasculature regenerated fully from auxin application, while *tmk1-1*, *tint6*, and *tint6 tmk1-1* mutants regenerated partially. The mutants *tint7* and *tint7 tmk1-1* showed no regeneration. White arrowheads indicate the wounding site. Black arrows indicate the regenerated vasculature from the applied source of auxin. Scale bar, 100 µm.

(b) Wounding of the stem of *DR5rev::GFP* and *PIN1::PIN1-GFP* triggered the complete development of auxin and PIN1 channels respectively (indicated by white arrows), but not in *tint5*, or *tint7* mutant. 4 DAW. Scale bar, 100 µm.

(c) Quantification of vasculature formation after wounding in *DR5rev::GFP* and *PIN1::PIN1-GFP* lines, categorized as complete (fully formed channels), partial (limited or partially developed channels), and none (no channels formed) at 4 DAW. n=10 plants per genotype.

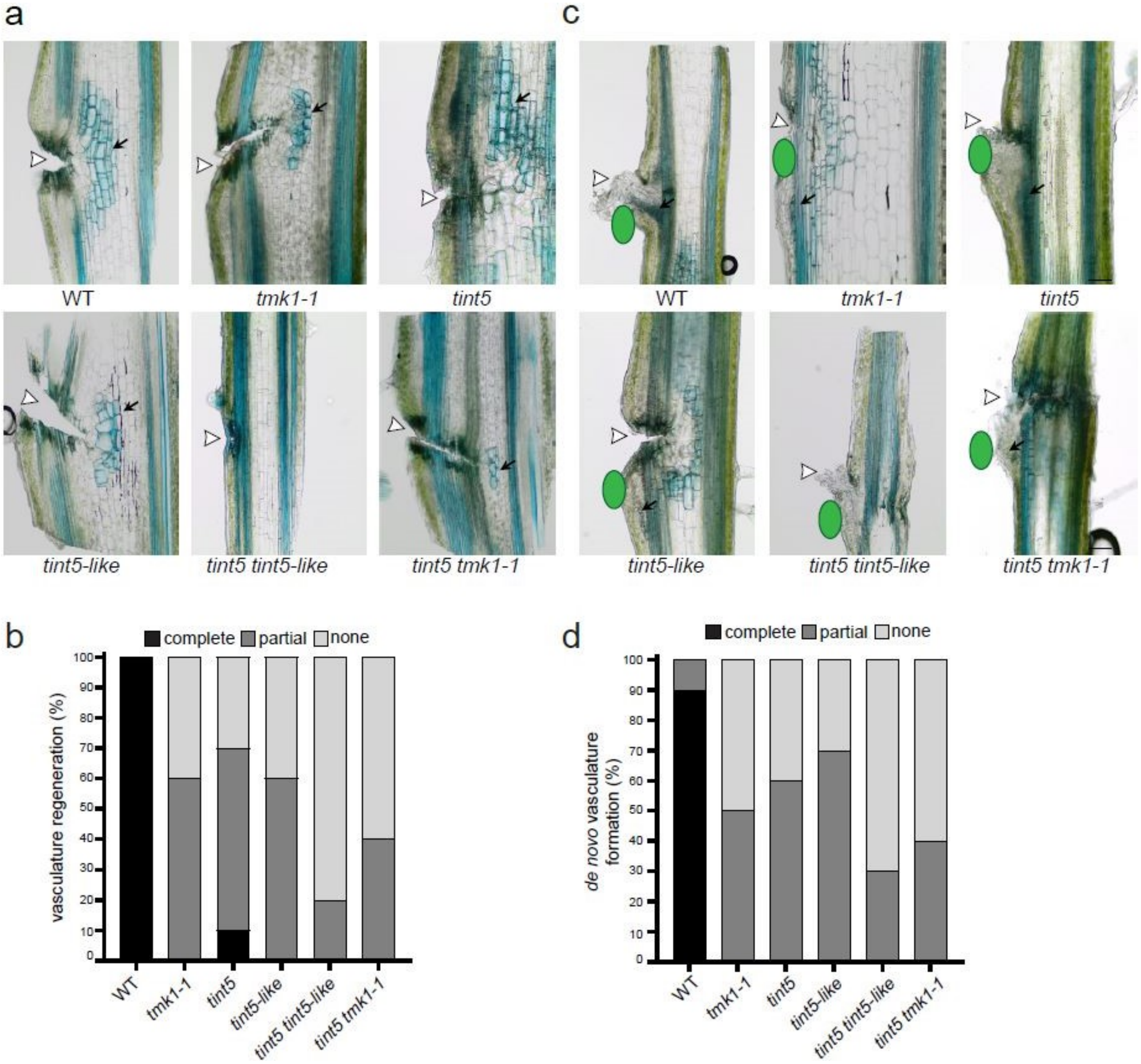

**Figure S7- TINT5 and TINT5-like in canalization and vasculature regeneration after wounding**

(a) Representative images of the TBO staining at 6 DAW showing the vasculature regeneration in the inflorescence stems. WT vasculature regenerated completely around the wound, while *tmk1-1*, *tint5*, *tint5-like*, and *tint5 tint5-like* showed defects in regeneration. The double mutant *tint5 tmk1-1* developed only single vessels. White arrowheads indicate the wounding site. Black arrows indicate the regenerated vasculature. Scale bar, 100 µm.

(b) Quantification of the vasculature regeneration in *tmk1-1*, *tint5*, *tint5-like* single and double mutants, categorized as complete (fully formed vasculature), partial (limited and partially developed vasculature), or none (no vessel formed around the wound) at 6 DAW. n=10 plants per genotype.

(c) Representative images of the TBO staining at 6 DAA of *de novo* vasculature formation and canalization from exogenous application of auxin (green ovals) following wounding. WT vasculature regenerated fully from auxin application, while all the shown mutants showed defects in regeneration. White arrowheads indicate the wounding site. Black arrows indicate the regenerated vasculature from the applied source of auxin.

(d) Quantification of *de novo* vasculature formation from a local source of auxin in *tmk1-1*, *tint5*, *tint5-like* single and double mutants, categorized as complete (fully formed vasculature), partial, and none (no channels formed) at 6 DAA. n=10 plants per genotype.

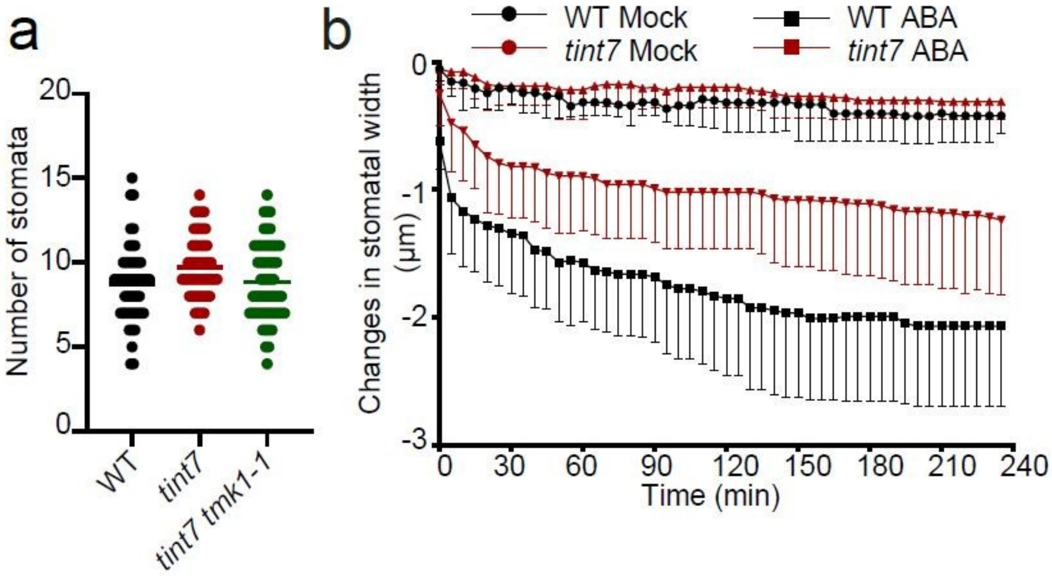

**Figure S8- TINT7 function in regulating stomatal movement**

(a) Graph showing the number of stomata in a given area of cotyledons from WT, *tint7*, and *tint7 tmk1-1*. The mutants exhibited a similar number of stomata compared to WT. n ≥ 50 per genotype.

(b) Quantification of stomatal closure kinetics after ABA treatment. The changes in stomatal width were measured every 5 minutes, and the average change with SD is displayed in the graph. In mock conditions, stomatal width in WT and *tint7* remained largely unchanged. However, when treated with ABA, *tint7* stomata closed less than WT. n ≥ 9 per genotype.

**Supplementary table**

| | |
|---|---|
| **For transgenic lines pUBQ10** | |
| attB1TINT1CDS-Fw | GGGGACAAGTTTGTACAAAAAAGCAGGCTATGAAGAACAAGACCAATTTAG |
| attB2TINT1CDS-Rv | GGGGACCACTTTGTACAAGAAAGCTGGGTcCAATCGGACAAAGGACCT |
| attB1TINT2CDS-Fw | GGGGACAAGTTTGTACAAAAAAGCAGGCTATGGAAGCTTTGAGGATTTA |
| attB2TINT2CDS-Rv | GGGGACCACTTTGTACAAGAAAGCTGGGTcCAATCGGACAAAGGACCT |
| attB1TINT3CDS-Fw | GGGGACAAGTTTGTACAAAAAAGCAGGCTATGACCCGTGATGACAAATT |
| attB2TINT3CDS-Rv | GGGGACCACTTTGTACAAGAAAGCTGGGTTTTCATGAGTGAGAACTCCT |
| attB1TINT4CDS-Fw | GGGGACAAGTTTGTACAAAAAAGCAGGCTATGAGTAGAGGAAGATCTTT |
| attB2TINT4CDS-Rv | GGGGACCACTTTGTACAAGAAAGCTGGGTcCAGTCTCTCTCAATCTCTT |
| attB1TINT5CDS-Fw | GGGGACAAGTTTGTACAAAAAAGCAGGCTATGCAGATTTCTCTTCAAATC |
| attB2TINT5CDS-Rv | GGGGACCACTTTGTACAAGAAAGCTGGGTcCATTCATATATTGCTTTCA |
| TINT6CDS-Fw | ggccgccccccttcaccATGGAGATTTCTTTGATGAA |
| TINT6CDS-Rv | cggcgcgcccacccttTAATCGTGGTCCAGAGAGCTCAAT |
| TINT7CDS-Fw | ggccgccccccttcaccATGAGAGGGAGGATATGTTTCTTG |
| TINT7CDS-Rv | cggcgcgcccacccttTAAAATTTCTTTGGAGGAGTTTGCTT |
| **For transgenic lines pTINT** | |
| pTINT5-Fw | ttgtatagaaaagttgggcAGCATTCACAAGGAATAAGCA |
| pTINT5-Rv | actttttgtacaaaattgtAGTGAAGATGTGTTTTGCCTT |
| pTINT5like-Fw | ttgtatagaaaagttgggcTGACACTCATTTCTCACCGAC |
| pTINT5like-Rv | actttttgtacaaaattgtTCTCTCAGGAAGAATTACAGAGA |
| TINT5CDS-Fw | ggccgccccccttcaccATGCAGATTTCTCTTCAAATCCATT |
| TINT5CDS-Rv | cggcgcgcccacccttCCATTCATATATTGCTTTCATGGATGATT |

| | |
|---|---|
| TINT5likeCDS-Fw | ggccgccccccttcaccATGCATAGTTCCTCTAAAAGCCAG |
| TINT5likeCDS-Rv | cggcgcgcccacccttCAATAGTTCTGAACCACCAAGCCC |
| **For GUS lines** | |
| pTINT6-Fw | ttgtatagaaaagttgggcATTAAGGCGGTTGGTGAC |
| pTINT6-Rv | acttttttgtacaaaattgtCGTAGAAAAAGTATAAGAATTCTCT |
| pTINT7-Fw | ttgtatagaaaagttgggcATCAACTTCCGTCTTACATGCAT |
| pTINT7-Rv | acttttttgtacaaaattgtTTTTACAAGTTAGCTTTGGTAAGAACAC |
| **For genotyping** | |
| LBb1.3 | ATTTTGCCGATTTCGGAAC |
| tint1 SALK053366_LP | AGAGAAACGAAGGAATCTCGC |
| tint1 SALK053366_RP | ATGTTCCACATATCCGAAAGG |
| tint2 SALK033657_LP | TTTCTTTTCTCCACGGTGTTG |
| tint2 SALK033657_RP | AAGAAAAGATTCCAGTGCAAAAAC |
| tint3 SALK086592_LP | TTTGGATCTTCGTACAAAGCG |
| tint3 SALK086592_RP | ATGCGAATAAATAACCCTGGC |
| tint4 SALK094070_LP | AAACTGCCGAAAGCACATATG |
| tint4 SALK094070_RP | AAAAAGAAGGTTGGGGATGTG |
| tint5 SALK131589_LP | TTGGTTAGCGTTCTTGGACAC |

| | |
|---|---|
| tint5 SALK131589_ RP | CATGGATGATTCATGATTCCC |
| tint5-like SALK078409_ LP | CTAAAAGCCAGGCCTTTTCTC |
| tint5-like SALK078409_ RP | GATCACACCGATGATAATGCC |
| tint6 SALK123502_ LP | AACAAAATCGAACCTTGGAGG |
| tint6 SALK123502_ RP | GCAATGGTCAAGTTCGAAAAG |
| tint7 SALK101617_ LP | GGTATCTTCCCTTCAAGCTGG |
| tint7 SALK101617_ RP | TATCTCGCAAGCTGGAATCAG |
| tmk1-1_LP | CCAGTTCCTGCGTCTTTGTT |
| tmk1-1_RP | TTAACCGCAATCTTCGTTCC |

**Table S1. Primers used for genotyping and generating constructs**
